# Ancient and modern geochemical signatures in the 13,500-year sedimentary record of Lake Cadagno

**DOI:** 10.1101/2021.10.04.463023

**Authors:** Jasmine S. Berg, Mathilde Lepine, Emile Laymand, Xingguo Han, Hendrik Vogel, Marina A. Morlock, Niroshan Gajendra, Adrian Gilli, Stefano Bernasconi, Carsten J. Schubert, Guangyi Su, Mark A. Lever

## Abstract

Although lake sediments are globally important organic carbon sinks and therefore important habitats for deep microbial life, the deep lacustrine biosphere has thus far been little studied compared to its marine counterpart. To investigate the impact of the underexplored deep lacustrine biosphere on the sediment geochemical environment and vice versa, we performed a comprehensive microbiological and geochemical characterization of a sedimentary sequence from Lake Cadagno covering its entire environmental history since formation following glacial retreat. We found that both geochemical gradients and microbial community shifts across the ∼13.5 kyr subsurface sedimentary record reflect redox changes in the lake, going from oxic to anoxic and sulfidic. Most microbial activity occurs within the top 40 cm of sediment, where millimolar sulfate concentrations diffusing in from the bottom water are completely consumed. In deeper sediment layers, organic carbon remineralization is much slower but microorganisms nonetheless subsist on fermentation, sulfur cycling, metal reduction, and methanogenesis. The most surprising finding was the presence of a deep, oxidizing groundwater source. This water source generates an inverse redox gradient at the bottom of the sedimentary sequence and could contribute to the remineralization of organic matter sequestered in the energy-limited deep subsurface.

## INTRODUCTION

Globally lake sediments accumulate organic carbon at an estimated annual rate of about 42 Tg yr^-1^ (Dean and Gorham, 1998), representing a significant sink for organic matter and a huge potential reservoir for microbial life. Continued sedimentation over geologic time buries deposited organic matter, including microbial cells, along with inorganic minerals and pore water deeper and deeper, thus preserving geochemical signatures from the time of deposition. Furthermore, due to the great sensitivity of lakes to environmental changes on the surrounding land (e.g. glacial retreat, permafrost thawing, soil development), lake sediments can serve as natural archives of regional climatological and ecological variations over time.

The influence of microorganisms on the sedimentary carbon reservoir is greatest in surface sediments, where much of the deposited organic matter is microbially accessible and easily degradable, and high energy microbial electron acceptors, such as O_2_ and nitrate are present (Carlton et al., 1989). Yet, even in deeper layers, in which these electron acceptors are depleted and the residual organic matter is increasingly recalcitrant to degradation, microbial activity can have a significant impact on the storage of organic carbon over time scales of thousands to millions of years. In these deeper, anoxic sediments, the microbial breakdown of organic carbon is carried out by a network of microorganisms. Primary fermenters perform the initial extracellular hydrolysis of organic macromolecules and gain energy by converting the released mono- and oligomers to smaller compounds, such as H_2_, short-chain organic acids, or alcohols (e.g., (Schink, 1997)). The resulting smaller compounds are then terminally oxidized to CO_2_ or CH_4_ by respiring microorganisms, using metal oxides, sulfate, or CO_2_ as electron acceptors. Alternatively, short-chain organic acids and alcohols produced by primary fermenters may first be converted to H_2_ or C1 compounds by secondary fermenters before being terminally oxidized by respiring microorganisms to CO_2_ and/or CH_4_.

In anoxic lake sediments with high sedimentation rates, CO_2_ reduction to methane (methanogenesis) and iron (Fe(III)) reduction are the dominant microbial respiration reactions (Lovley and Phillips, 1986; Capone and Kiene, 1988; Roden and Wetzel, 1996). Despite generally low sulfate concentrations in freshwater environments, efficient recycling of this terminal electron acceptor may increase the importance of sulfur respiration pathways (Bak and Pfennig, 1991; Urban et al., 1994; Hansel et al., 2015). While evidence shows that these biogeochemical processes continue to be important in deep submarine sediments (Onstott et al., 1999), little data is available from deep lake sediments. Differences in trophic state, water chemistry, and the quality and quantity of organic matter input between lakes and oceans make it difficult to generalize across such contrasting environments. Moreover, it is known from marine sediments that metabolic activities may deviate from the standard energetic model and that deep geologic interfaces, e.g. of sediments with underlying aquifers, can supply dissolved electron acceptors, such as O_2_, nitrate, or sulfate to deep sediment layers (D’Hondt et al., 2004; Parkes et al., 2005; Jørgensen et al., 2020).

The microorganisms driving biogeochemical processes in deep lake sediments have been much less studied than their marine counterparts. Since microbial activity and cell concentrations in subseafloor sediments correlate with the sedimentation rate of organic matter (Kallmeyer et al., 2012), the lacustrine subsurface biosphere can also be expected to vary greatly across different lake environments. At this time, deep sediment microbial communities of only two lakes have been investigated (Vuillemin and Ariztegui, 2013; Vuillemin et al., 2018; Thomas et al., 2020) and far more studies will be needed to understand lake subsurface ecosystems. Thus far, microbial diversity in these lacustrine subsurface environments has proven surprisingly similar to that in deep marine sediment, despite differences in initial seed communities (Vuillemin et al., 2018; Thomas et al., 2020).

Due to the sequential changes in its redox and sediment depositional history, Lake Cadagno in the Piora Valley of the Swiss Alps provides a suitable limnological record for studying biogeochemical processes as a function of sediment age and lithology. Situated at 1921 m altitude, this lake formed during glacial retreat and has since undergone transformations from oxic, to suboxic, to completely anoxic bottom waters (Wirth et al., 2013). Today the permanently stratified lake is often cited as an Archaean ocean analogue for its sulfate-rich (∼2 mM), anoxic bottom water, which harbors biogeochemical processes that may have been widespread on early Earth (Canfield, 1998; Poulton et al., 2004). In order to understand how environmental changes alter the (microbial) cycling of carbon, sulfur, and metals (Fe, Mn), and microbial community compositions through time, we analyzed the complete sedimentary record of Lake Cadagno. By combining pore water and solid-phase geochemical analyses with quantitative and high-throughput sequencing of 16S rRNA genes, we reveal an unexpected distribution of biogeochemical processes and microbial communities influenced by a subsurface aquifer.

## METHODS

### Study site and field sampling

Lake Cadagno is a crenogenic meromictic lake located in the Swiss Alps at 1921 m above sea level. Subaquatic springs flowing through dolomitic bedrock supply the lake with high concentrations of Mg^2+^, Ca^2+^, HCO_3_^-^ and SO_4_^2-^, the latter of which is respired by sulfate-reducing microorganisms generating a sulfidic hypolimnion and underlying sediments. A previous study of a ∼10-m sedimentary succession reaching from the sediment-water interface to the underlying late glacial sediments revealed that the 21 m-deep lake basin formed after glacial retreat about 12.5 thousand years ago and underwent a several-hundred-year redox transition interval before shifting to fully anoxic, sulfidic conditions (Wirth et al., 2013). Correspondingly, the sedimentary record hosts three main lithological units: late glacial deposits poor in organic matter, a Mn-enriched redox transition zone, and metal sulfide-rich sediments. Lipid biomarkers of anoxygenic phototrophic bacteria have been recovered from the metal sulfide-rich layers and indicate anoxic, hydrogen sulfide-rich conditions in overlying lake water throughout the time of deposition (Wirth et al., 2013). These euxinic sediments are additionally characterized by discrete sediment layers of different origins: laminated to thinly bedded pelagic lacustrine muds, graded coarse-grained terrestrial flood deposits, and mass-movement deposits containing remobilized pelagic and/or terrestrial sediment. To investigate how the depositional history, lithology, and redox history of these sediments have shaped biogeochemical processes and associated microbial communities through time, a second piston-coring campaign was undertaken in Lake Cadagno in August 2019.

Three sets of piston cores (Supplementary Fig. 1) were recovered using a Uwitec coring platform (Uwitec, AT) of the ETH Zurich’s Institute for Limnogeology. This coring platform recovers 3-m long and 6-cm diameter core sections using a percussion piston-coring system. The three piston cores were obtained from parallel boreholes in the deepest part of the lake (8.71201 E and 46.55060 N, no more than +/- 0.00003 decimal degrees or ∼3 meters apart). Florist foam soaked with lake surface water was used to fill any remaining empty space in the top of the core liner before sealing with rubber caps at each end. Each core section was carried to the Alpine Biology Center field laboratory on shore for immediate sample processing. For high-resolution analyses of undisturbed surface sediments, additional short cores were retrieved from the same location using a UWITEC gravity corer using clear plastic liners with 1 m length and 9 cm inner diameter.

The first set of long cores was reserved for non-destructive imaging analyses. The second set of long cores was dedicated to pore water extraction using syringes connected to Rhizons (0.2 μm pore size, Rhizosphere) inserted into holes drilled horizontally every 10 cm into the core liners after collection. Rhizons, stopcocks and syringes were first flushed with 2– 3 mL of pore water to remove contaminant air. Porewater was then distributed into separate vials with appropriate fixatives for downstream analyses. Samples from the top 30 cm of each core were discarded due to the likely infiltration of surface water in the florist foam. The third set of long cores was dedicated to analyses of DNA, dissolved gases, solid-phase C, Fe, and S pools, and physical properties. Sampling windows were cut into the core liners every 20 cm using a hand-held vibrating saw and potentially contaminated sediment in contact with the liner was scraped away. Samples were then taken using sterile, cut-off syringes. After sampling, all cores were sliced into 1-m sections and split longitudinally for high-resolution photography to enable alignment of parallel cores and to establish a continuous composite core record.

### High-resolution imaging and age model construction

The chronology of the 2019 core composite is based on 9 radiocarbon dates which were transferred from the previously studied 2009 Lake Cadagno core succession (Wirth et al., 2013). Transfer of dates is based on aligning the characteristic lithologies from which the dates were obtained between the two composite sediment core successions. Upon age-depth modeling using the Bayesian statistics Bacon.R software (Blaauw and Christen, 2011) ^14^C ages were converted into calibrated ^14^C ages (cal. a BP) using the IntCal13 calibration curve (Reimer et al., 2013). We removed event deposits (flood layers, slumps) >2 mm prior to age-depth modelling and reinserted these into the chronostratigraphy following age-depth modeling using a constant age for each individual deposit. The age-depth model was not extended into the basal coarse-grained late glacial deposits due to the obvious differences in depositional environment under which these were deposited and lack of reliable dateable material in these primarily rapidly deposited clastic sediments.

### Porewater Chemistry

Samples for analysis of dissolved metals (Fe, Mn) were acidified with 5 µl of 30% HCl per 2 mL to prevent precipitation and measured by inductively coupled plasma optical emission spectroscopy (ICP-OES, Agilent Technologies 5100). Porewater for dissolved inorganic carbon (DIC) analysis was filled into 1.5 mL borosilicate vials and capped without headspace to avoid degassing of CO_2_ and then stored at 4ºC for up to 8 weeks. Samples were transferred to He-flushed Exetainers immediately prior to addition of 85% phosphoric acid and analysis on a gas chromatograph (GasBench II, Thermo Fisher Scientific) coupled with a mass spectrometer (Delta V, Thermo Fisher Scientific). For dissolved sulfide analyses, pore water was fixed with Zn-acetate solution to 0.5% final concentration and quantified photometrically using the methylene blue method (Cline, 1969). Samples for dissolved ion (PO_4_^3-^, NO_2_^-^, NO_3_^-^, SO_4_^2-^, NH_4_^+^) analyses were immediately frozen at -20°C until analysis on an ion chromatograph (DX-ICS-1000, DIONEX) equipped with an AS11-HC column. Carbonate buffer (3.2 mmol l^-1^ Na_2_CO_3_ and 1 mmol l^-1^ NaHCO_3_) was used as eluent at a flow rate of 1 ml min^-1^ with a total run duration of 14 min. Nitrite, nitrate, phosphate and sulfate eluted at 5.8, 8.0, 9.7, and 11.9 min, respectively. Ammonium was determined photometrically using the indophenol blue method (Kempers and Kok, 1989).

### Dissolved gases

For dissolved methane quantification, 3 cm^3^ of sediment was immediately transferred to 20 mL glass vials containing 7 mL of 10% NaOH, sealed with a butyl rubber stopper, and homogenized by shaking (Blees et al., 2014). Methane concentrations in the headspace were measured using a gas chromatograph (GC, Agilent 6890N) with a flame ionization detector and He as a carrier gas.

### Solid Phase Carbon, Nitrogen, Iron, and Sulfur Analyses

Using a cut-off plastic syringe, 5 cm^3^ of sediment were obtained from each depth interval, transferred to clean glass vials with screw caps, and stored at -20°C until further processing. Porosity was determined from weight loss after heating the sediment at 70°C until complete dryness. Total carbon (TC) and total nitrogen (TN) were determined from this dried sediment by EA-IRMS as described below. Total organic carbon (TOC) was determined after acid-extraction of inorganic carbon with concentrated 6 N HCl. Total inorganic carbon (TIC) was calculated as the difference between TC and TOC.

An additional ∼10 cm^3^ (or more) of fresh sediment from each depth were transferred to sterile, gas-tight plastic bags (Whirl-Pak) and sealed after pressing out all the air before freezing at -20°C. For solid phase iron and sulfur extractions, bagged sediments were thawed in a cold-water bath and then, avoiding the sediment in contact with the plastic bag, a sub-sample was quickly transferred to a degassed glass vial and freeze-dried overnight. Less than 100 mg of the dried sediment was weighed into a 15 mL Falcon tube containing 0.25 mM HCl for reactive iron analysis, and the remainder of sediment was stored anoxically for sulfur extractions. Reactive iron was extracted for 1 h on a shaker prior to centrifugation and photometric determination of Fe(II)/Fe(III) in the supernatant using the ferrozine assay (Stookey, 1970).

Elemental sulfur was extracted three times under an N_2_ atmosphere with degassed 100% methanol. During each step the methanol-sample mixture was sonicated for 10 min in an ice bath, centrifuged, and then the methanol was pipetted into a clean vial. Methanol extracts were analyzed by ultrahigh pressure liquid chromatography (UPLC) using a Waters Acquity H-class instrument with an Aquity UPLC BEH C18, 1.7 µm, 2.1 × 50 mm column (Waters, Japan) and a PDA detector (absorbance wavelength set to 265 nm). The injection volume was 10 µl with methanol as eluent flowing at 0.2 ml min^-1^. Elemental sulfur eluted at 4.14 min.

### Concentration and Stable Isotopic Analyses of Carbon and Nitrogen Pools

The concentrations and isotopic compositions of nitrogen and carbon in different sediment fractions were determined simultaneously using a Flash-EA 1112 (ThermoFisher Scientific) coupled to an isotope ratio mass spectrometer (IRMS, Delta V, ThermoFisher Scientific). Isotope ratios are reported in the conventional -notation with respect to atmospheric N_2_ (AIR) and Vienna Pee Dee Belemnite (V-PDB) standards, for nitrogen and carbon, respectively. The system was calibrated with IAEA-N1 (^15^N = +0.45), IAEA-N2 (^15^N = +20.41) and IAEA N3 (^15^N = +4.72) reference materials for nitrogen, and NBS22 (^13^C =-30.03) and IAEA CH-6 (^13^C =-10.46) for carbon. The reproducibility of measurements was better than 0.2‰ for both elements.

### Diffusive flux calculations

Fluxes of SO_4_^2-^, and CH_4_ diffusing into the upper and lower sulfate-depletion zones were calculated from pore water concentration gradients according to Fick’s first law:

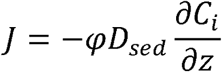

∂X where *J* is the diffusive flux (in mmol m^-2^ d^-1^), *φ* is the mean local sediment porosity, *D*_sed_ is the sediment diffusion coefficient (in m^2^ s^-1^) for *in situ* conditions and corrected for tortuosity (Boudreau, 1997), *C* is the concentration of molecule *i* (in mmol per m^3^ of pore water), and *z* is the sediment depth (in mblf). Molecular diffusion coefficients (D_0_) selected for the *in situ* temperature of Lake Cadagno sediments were 1.012 × 10^−9^ m^2^ s^-1^ and 5.808 × 10^−10^ m^2^ s^-1^ for SO_4_^2-^ and CH_4_, respectively (as in (Su et al., 2019)).

### DNA extraction & sequencing

For DNA analyses, ∼4 cm^3^ of sediment from each depth were transferred to sterile 5-cm^3^ cryovials, immediately flash-frozen in liquid N_2_, and stored at -80°C until further processing. DNA was extracted according to (Lever et al., 2015) using Lysis Protocol II followed by 2× washing with chloroform-isoamylalcohol (24:1), precipitation with ethanol-sodium chloride-linear polyacrylamide, and post-extraction clean up with the Norgen CleanAll DNA/RNA Clean-Up and Concentration Micro Kit (further details in (Lever et al., 2015)). Prior to lysis, 0.2 g sediment aliquots were soaked in 100 µL hexametaphosphate solution to minimize DNA adsorption to sediment grains. The concentration of hexametaphosphate was 10 mM in all samples except those from organic-poor glacial clay (≥782 cmblf), where they were increased to 100 mM due to the otherwise very high adsorptive losses.

16S rRNA gene concentrations in purified genomic DNA extracts were quantified using SYBR Green I Master on a LightCycler 480 II (Roche Molecular Systems, Inc.). The primer pairs for Bacteria and Archaea were Bac908F_mod (5’
s-AAC TCA AAK GAA TTG ACG GG-3′) (Lever et al., 2015)/ Bac1075R (5′-CAC GAG CTG ACG ACA RCC-3′) (Ohkuma and Kudo, 1998) and Arch915F_mod (5′-AAT TGG CGG GGG AGC AC-3′) (Cadillo Quiroz et al., 2006)/Arch1059R (5′-GCC ATG CAC CWC CTC T-3′) (Yu et al., 2005), respectively. The thermal cycler protocols followed those outlined in Lever et al., (2015).

For amplicon sequencing, the V3-V4 hypervariable regions of bacterial and archaeal 16S rRNA genes was amplified using the primer pairs S-D-Bact-0341-b-S-17 (5′-CCT ACG GGN GGC WGC AG-3′)/S-D-Bact-0785-a-A-21 (5′-GAC TAC HVG GGT ATC TAA TCC-3′) (Herlemann et al., 2011) for Bacteria and S-D-Arch-0519-a-A-19 (5′-CAG CMG CCG CGG TAA HAC C-3′) (Sørensen and Teske, 2006)/967Rmod (5′-GTG CTC CCC CGC CAA TT-3′) (Cadillo Quiroz et al., 2006) for Archaea. Additionally, to investigate potential primer biases, 16S rRNA genes from a subset of samples spanning the entire sediment column were amplified using the universal prokaryotic primers Univ519F (5’
s-CAG CMG CCG CGG TAA-3′) / Univ802R (5′-TAC NVG GGT ATC TAA TCC-3′) (Claesson et al., 2009; Wang and Qian, 2009). Amplicon libraries were prepared in-house starting with a booster PCR (threshold number of qPCR cycle plus three additional cycles) to obtain similar amplicon concentrations across all samples, followed by a tailed-primer PCR (10 cycles) and an index PCR (8 cycles). Further details can be found in (Deng et al., 2020) for the universal bacterial and archaeal primer pairs, and in Han et al. (2020) for the universal primer pair). Paired-end sequencing was performed on a MiSeq Personal Sequencer (Illumina Inc., San Diego, California, USA).

### Sequencing data processing and analysis

Phylogenetic analyses of DNA sequences were performed as described previously (Deng et al., 2020). For low biomass samples, potential contaminants were manually removed based on lists of common laboratory contaminants (Sheik et al., 2018; Karstens et al., 2019) and sequences present in negative controls (Supplementary Fig. 2). After decontamination 2,053,665 bacterial and 4,506,390 archaeal reads remained (details provided in supplemental files 1&2). Statistical analyses were performed in R v3.4.0 (http://R-project.org) using R Studio v1.1.442 (http://rstudio.com). Raw sequences have been deposited in Sequence Read Archive with accession numbers SAMN20601530-SAMN20601580 under the project PRJNA752598.

## RESULTS

### Lithological description and age model for Lake Cadagno sediment cores

The 9.6-m sediment succession recovered from Lake Cadagno spans the entire sedimentary history of the lake down to sediments that were deposited following glacial retreat from the depression (Fig. 1). The lithostratigraphy is characterized by alternating laminated to thinly bedded pelagic lacustrine sediment and turbidites of varying thickness (< 2-30 mm), originating from flood and mass-movement events) in the upper 778 cm. Turbidite deposits are mainly composed of coarse-grained terrestrially sourced materials with minor contribution from remobilized lacustrine sediment. Below 778 cm, predominantly detrital clastic late glacial deposits appear light gray and individual sediment beds are delimited by contrasting grain-sizes ranging from clay to sand (Supplementary Fig. 1; (Wirth et al., 2013). The transition from late glacial to lacustrine deposits is marked by a 20-cm thick succession of Mn-oxide rich layers indicating the transition to seasonal stratification. The lacustrine sediment succession from 778 cm to the sediment-water interface spans the past ∼9.95 kyrs continuously based on the available radiocarbon dates and age-depth modelling that also takes upcore sedimentation rates into account (Fig. 1). Based on this age-model we estimate the duration of the transition period with Mn-oxide layer deposition to have lasted between ∼9.95 (780 cm) to ∼9.01 (760 cm) kyrs BP. Euxinic conditions persisted continuously since 9.01 (760 cm) kyr BP.

**Figure 1.**
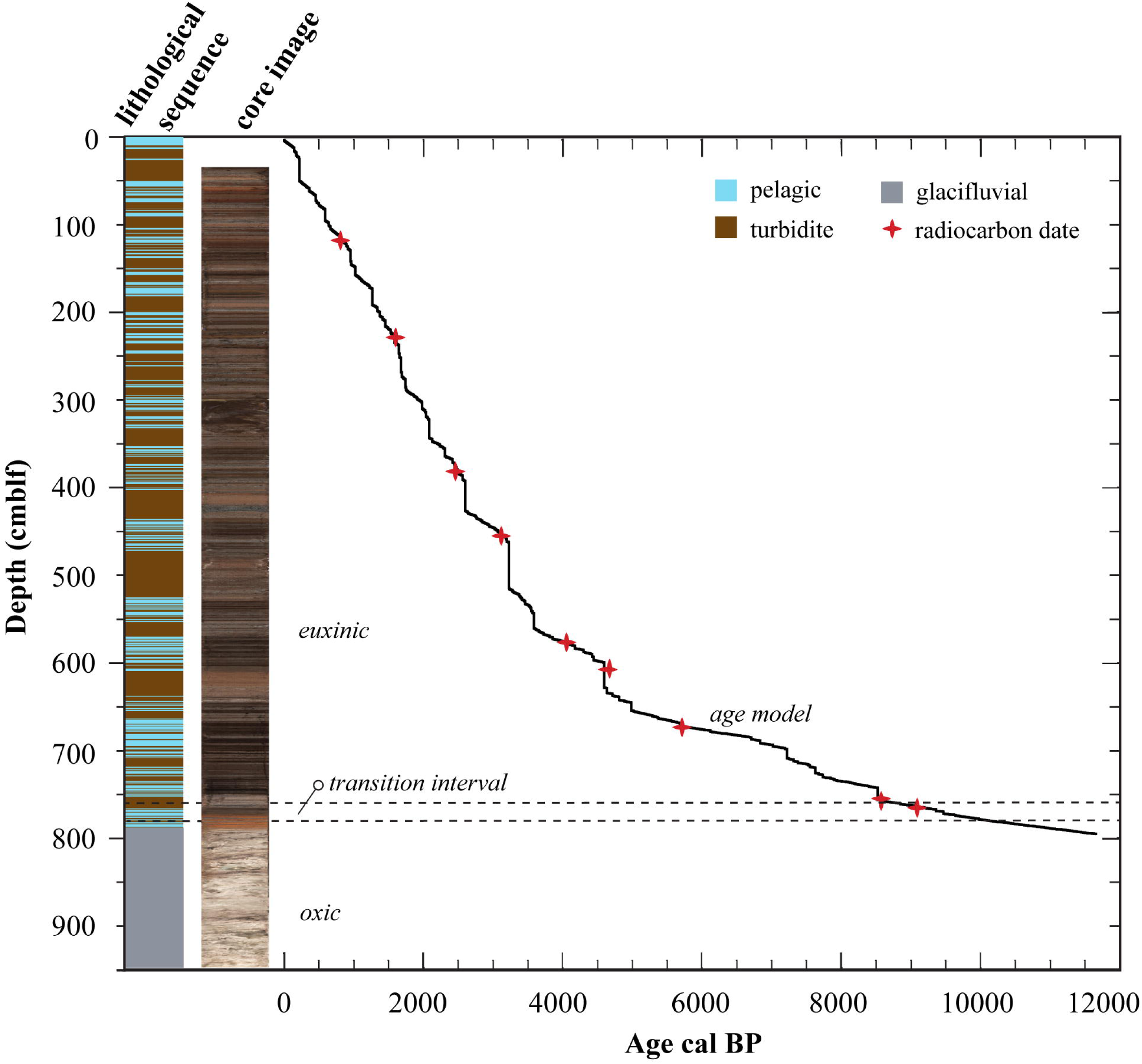
Lithological profile determined from a composite core image of the sedimentary sequence retrieved from Lake Cadagno. The age model was calibrated with previously obtained radiocarbon dates from corresponding sedimentary layers ((Wirth et al., 2013)).

### Strong geochemical gradients in surface sediments of Lake Cadagno

High-resolution geochemical profiles of the top 40 cm reveal strong vertical gradients of both solid and dissolved compounds in surface sediments. The TOC content is 18 wt % at the surface and drops rapidly to < 5% within the top 20 cm (Fig. 2A). The TIC is comparatively low and only a few values in the surface sediment are above the 0.8 % reproducibility limit of the instrument, with a maximum of 2.0 wt % TIC in the top 0-1 cm (Supplementary Fig. 3A). TN decreases in parallel with TOC, from 1.5 wt % at the surface to 0.2 wt % at 20 cm. Over the same interval, C:N ratios increase slightly, from 10.6 at the sediment surface to 11.7 at 20 cm. ^13^C-TOC values are lowest in surface sediment (□^13^C = -35 ‰) and become progressively higher with depth (^13^C = -27 ‰; Fig. 2B).

**Figure 2.**
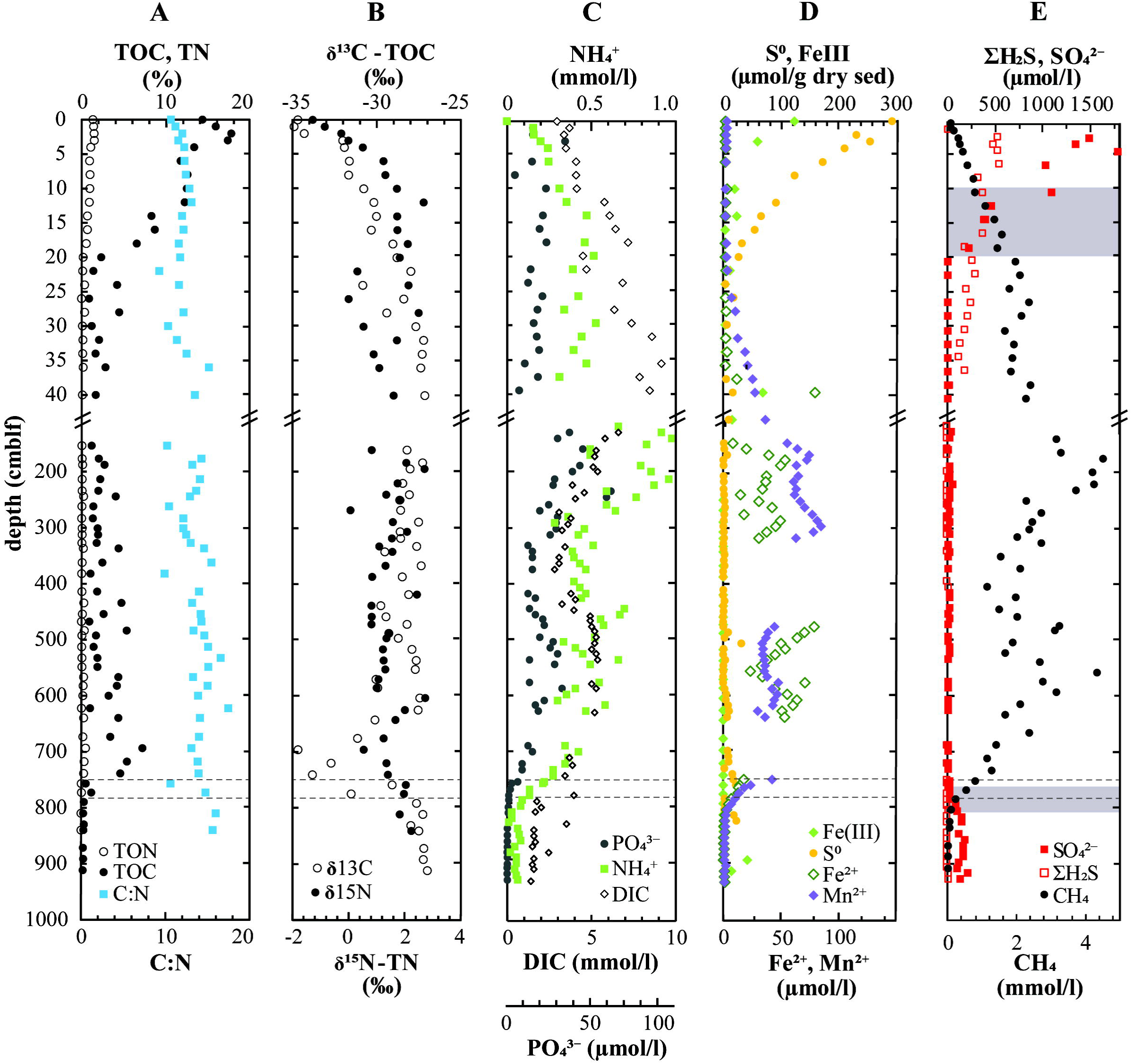
Geochemical profiles from the sediments of Lake Cadagno. Note the break in the y-axis showing the upper 40-cm of sediment in high-resolution. Dashed lines indicate the geological transition intervals and shaded regions in (E) denote the sulfate depletion zones. Large gaps in (D) are due to insufficient sample volume to measure all compounds.

The mineralization of organic matter is also evident from the increase in DIC and NH_4_^+^ concentrations with depth, from 2.9 mM and 150 µM at the surface to 9.1 mM and 540 µM at 40 cm depth, respectively (Fig. 2C). CH_4_ concentrations also increase near-linearly with depth, reaching ∼2.5 mM at 25 cm, and continuing to increase more gradually below (Fig. 2E). Neither NO_3_^-^ nor NO_2_^-^ were detected in surface sediments, but sulfate is depleted at approximately 40 cm depth without the accumulation of free sulfide (Fig. 2E). Due to the strong overlaps in CH_4_ and SO_4_^2-^ concentration profiles, we conclude that there is no true sulfate-methane transition zone. PO_4_^2-^ concentrations remain relatively constant between 100-200 µM (Fig. 2C) whereas dissolved Mn^2+^ and Fe^2+^ accumulate below 25 cm depth, (Fig. 2D).

Solid-phase analyses revealed up to ∼300 µmol S^0^/g dry sediment at the sediment surface. This S^0^ decreases steeply, stabilizing at low micromolar concentrations at 20 cm and below (Fig. 2D). There is a parallel decrease of reactive Fe-oxides within the top 5 cm of sediment which does not preclude the persistence of Mn-oxides or Fe-oxides that are not extracted with 0.25 mM HCl, and thus not considered reactive based on this method.

### Geochemical gradients across the 13.5 kyrs subsurface sedimentary record reflect historical redox changes in the lake

The TOC content of methanogenic sediments in the depth interval 160-778 cm fluctuates between 1.0-5.5% and TN between 0.1-0.4%, with no obvious depth-related trends, except for an overall increase in TOC that is most visible downcore of 500 cm (Fig. 2A). In parallel, there is a slight increase in C:N throughout this depth interval. Strong local differences can be attributed to sediment origin, with pelagic deposits being characterized by significantly (p < 0.01, Student t-test) lower C:N ratios (10.55 +/ 1.39) than mass movement deposits (13.75 +/- 1.68). The isotopic composition of TOC, but not TN, is also significantly lighter in lacustrine deposits (p < 0.01) than in turbidites.

There are relatively high concentrations of the reduced compounds Fe^2+^, Mn^2+^, and NH_4_^+^ as well as DIC in pore waters down to ∼670 cm. Methane exhibits a similar trend, reaching maximum concentrations of 4.5 mM (Fig. 2E), which is almost certainly an underestimate due to degassing evidenced by observed core expansion and bubble formation during sampling. Interestingly, the concentration minima of all of these compounds occur around 400-420 cm depth which corresponds to a thick mass-movement deposit (Fig. 1) also containing a local peak in TIC (1.8 wt %; Supplementary Fig. 3A). The low sulfate concentrations of 0-25 μM throughout this depth interval are possibly due to contamination of subsurface porewaters with sulfate-rich bottom waters during sediment coring.

Below 670 cm, porewater gradients of reduced metabolites (Fe, Mn, NH_4_^+^, and CH_4_) decrease sharply, reaching the limit of detection by 790 cm. Over the same depth interval, TOC (7.29 %) and TN (0.56 %) have local peaks at 670 cm that are followed by strong decreases to 0.36% TOC and < 0.02% TN at 790 cm. This decrease in TOC and TN contents coincides with a clear drop in the isotopic composition of TOC, from -27 to -30 ‰ in mid column sediment above 670 cm to values as low as -34.7 ‰ at 700 cm, followed by an increase back to ∼-28 ‰ at 790 cm and below (Fig. 2B). This local drop and subsequent increase in TOC isotopic values from 670 to 790 cm is accompanied by a slight gradual increase in TN isotopic values from ∼0.8 to 2.0 ‰. The distinct chemical and isotopic changes between 670-790 cm appear to be associated with a lithological transition at 760-790 cm, which corresponds to the anoxic-oxic transition period at 9.95-9.01 kyrs.

Late glacial sediments below 790 cm are characterized by extremely low TOC and TN content (Fig. 2A), reflecting the low productivity and terrestrial organic matter input. In these deep sediments, sulfate concentrations increase steeply from 4 µM at 778 cm to values of 100-210 µM at 820 cm and below (Fig. 2E). Active sulfate reduction is evidenced by a small peak in free sulfide (9 µM) around 810 cm, which likely reacts with Mn- or Fe-oxides there forming abundant (22 µM) zero-valent sulfur (Fig. 2D).

### Relationship of Bacterial and Archaeal abundances to sediment depth and organic carbon content

Bacterial and archaeal abundances based on 16S rRNA gene copy numbers are relatively stable in the top ∼30 cm and decrease below (Fig. 3A). Bacterial gene copies reach up to 7 × 10^8^ copies/g wet sediment, whereas archaeal gene abundances are an order of magnitude lower, with up to 8 × 10^7^ copies/g wet sediment. Below 30 cm, bacterial and archaeal gene copies only gradually decrease to 2 × 10^7^ and 1 × 10^7^ copies/g wet sediment, respectively, at 760 cm depth. There is only a very weak correlation between TOC content and total 16S gene copy numbers (Supplementary Fig. 4) indicating that other factors explain the variation in gene copy numbers in the mid-column sediment. Nonetheless, a peak in bacterial (8 × 10^7^ copies/g wet sediment) and archaeal (8.18 × 10^7^ copies/g wet sediment) gene abundances at 680 cm depth coincides with the region of higher TOC content (Fig. 2A) at the base of the euxinic mid-column sediment. Below this depth, i.e. throughout the lower sulfate depletion zone and underlying oxidized sediment, both bacterial and archaeal 16S rRNA gene copies drop drastically, to only several thousand copies in the deepest sediments. The Bacteria-to-Archaea 16S rRNA gene copy ratios (BAR) is stable around ∼8-12 in the top 20 cm, and then decreases gradually to values around ∼1 below 100 cm (Fig. 3B). BAR values then increase again locally in the lower sulfate depletion zone and underlying oxidized sediment.

**Figure 3.**
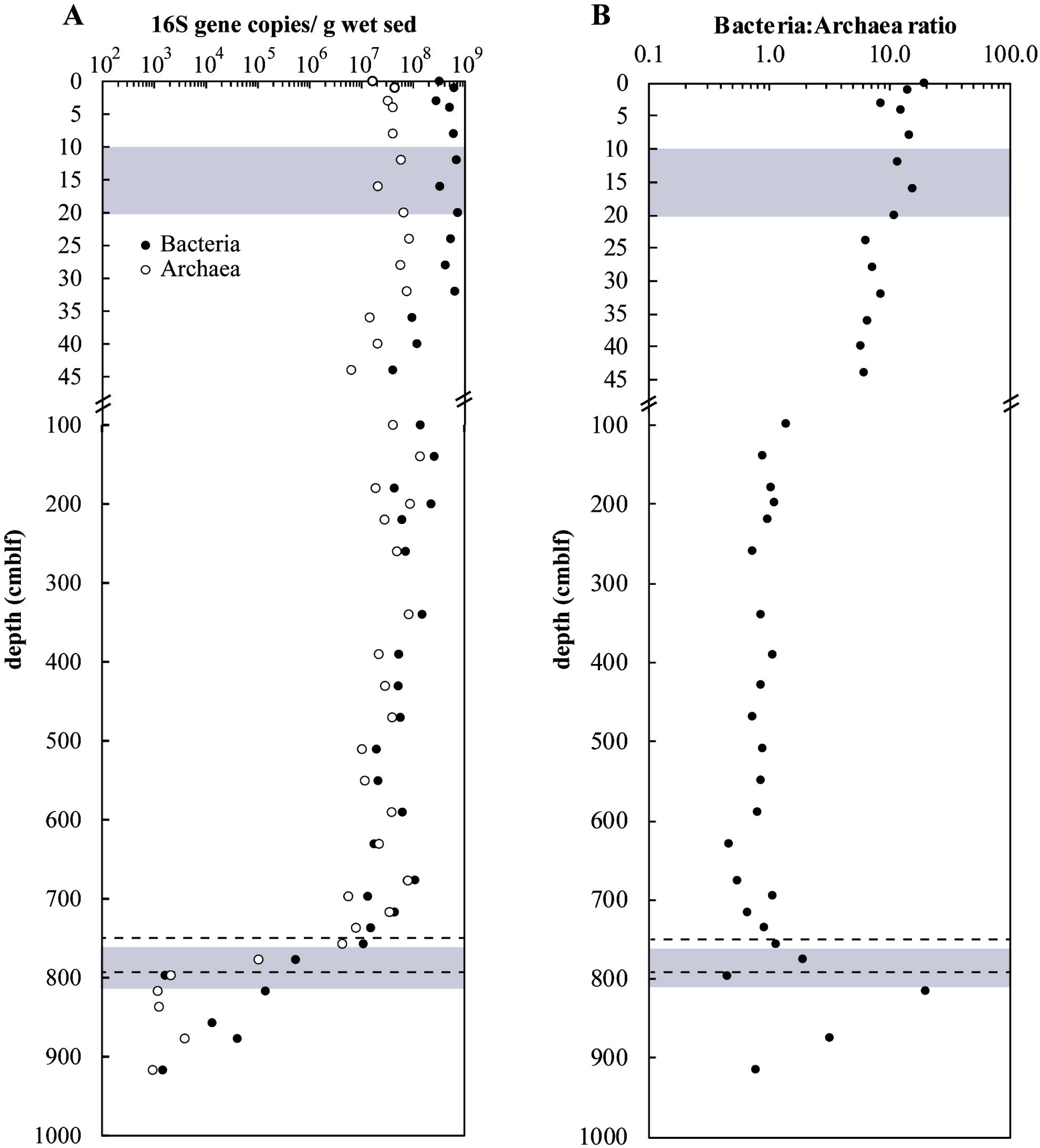
(A) Archaeal and bacterial 16S rRNA gene copy numbers and (B) BARs vs. depth in sediment of Lake Cadagno. Note the break in the y-axis showing the upper 45-cm of sediment in high resolution. Dashed lines denote the oxic-anoxic transition interval and shaded regions denote the sulfate depletion zones.

### Depositional history determines present-day microbial community zonation in Lake Cadagno

According to principal coordinates analysis of Bacterial and Archaeal diversity, microbial communities cluster into four distinct groups according to sediment age, sediment provenance, and/or redox conditions during sediment deposition (Fig. 4). These distinct groups can be classified into surface sediment (0-44 cm depth), euxinic sediment (150-760 cm), redox transition sediment from the deep oxic-anoxic transition (760-778 cm), and late glacial sediment (778-910 cm).

**Figure 4.**
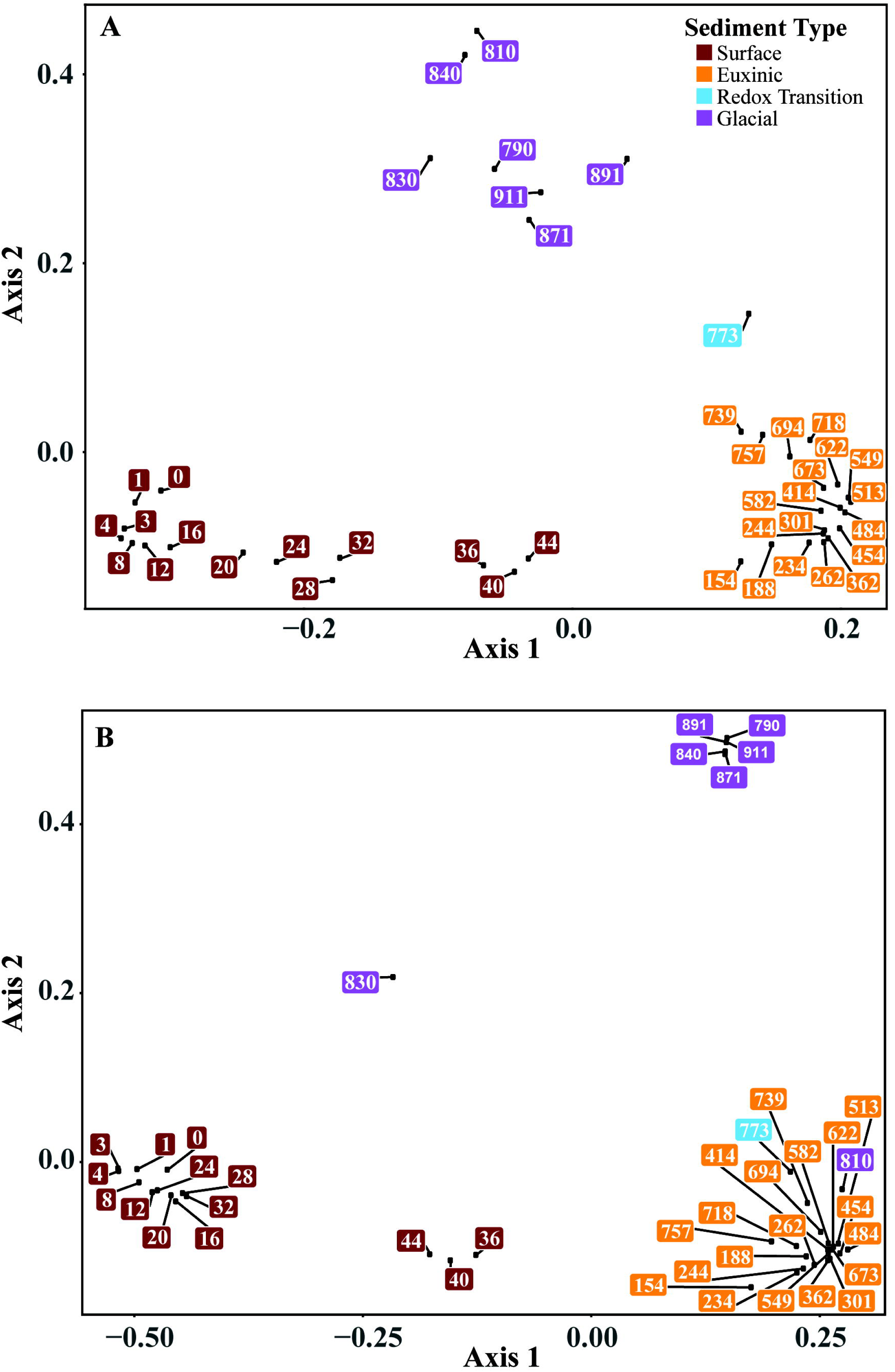
Principal coordinates analysis plot of (A) bacterial and (B) archaeal community composition versus lithological drivers. Sample names (depths) are color-coded according to sediment type.

These sediment groups, excluding the redox transition layer for which there is only one sample, are highly divergent from each other, sharing only 51 common Bacterial ZOTUs and 8 Archaeal ZOTUs. The transition in community structure appears to occur at different depths for Bacteria and Archaea. Bacterial populations from the redox transition sediment at 773 cm depth are intermediate but distinct from the strongly separated euxinic and late glacial sediments (Fig. 4A). In contrast, archaeal communities from the redox transition cluster with euxinic layers, both of which are clearly separated from late glacial sediments below 850 cm. Notably, the glacial sediment archaeal community at 810 cm clusters with the euxinic and redox transition layers, whereas the community at 830 cm appears distinct from all other sediment zones.

### Phylum- and class-level variations in bacterial communities

A total of 4,095 bacterial ZOTUs (2,053,665 reads) were recovered from the Lake Cadagno sediments. The vast majority, i.e. all but 358 bacterial ZOTUs, could be phylogenetically assigned at the phylum-level (Fig. 5). Of the 59 assigned bacterial phyla, only four were recovered from all sediment depths (Actinobacteria, Atribacterota, Firmicutes, and Proteobacteria).

**Figure 5.**
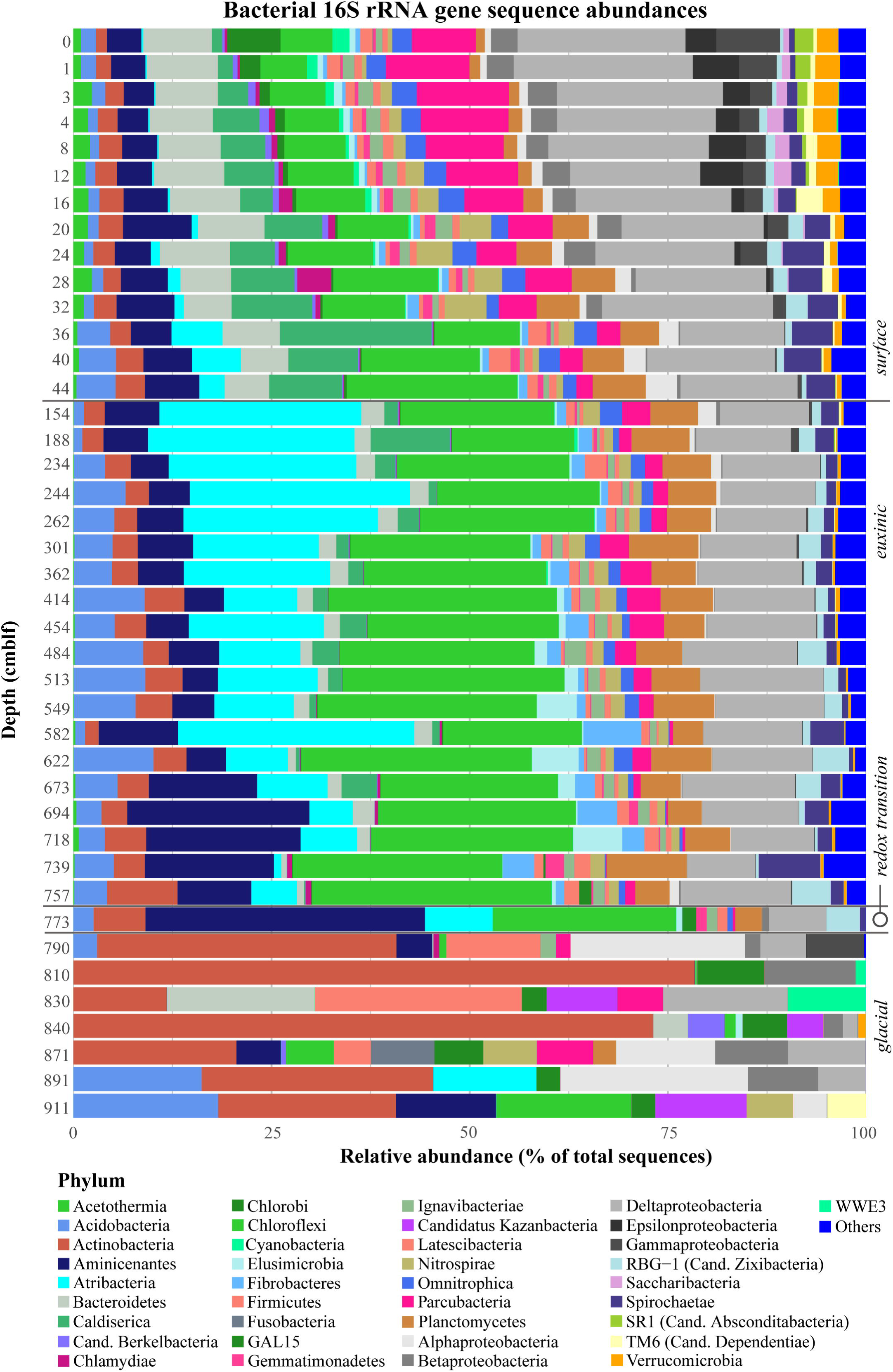
Relative abundances of bacterial 16S rRNA gene sequences with depth. Sediment geological transitions are indicated on the right.

Bacterial diversity is greatest (comprising 1781-2205 distinct ZOTUs) in the surface sediment although at least some of these microorganisms, i.e. the photosynthetic Cyanobacteria, Chromatiales, and Chlorobi, may simply constitute a planktonic fraction that has settled from the water column and is not metabolically active in sediments (Ravasi et al., 2012). The Proteobacteria, which are the dominant phylum in surface sediments, are comprised mainly of Beta-, Delta-, Gamma-, and Epsilonproteobacteria with low percentages of Alphaproteobacteria. The proportion of Proteobacteria decreases progressively from a total of 35 % in the top 0-1 cm to 19 % at 44 cm depth. Deltaproteobacteria represent 57-87% of the Proteobacteria in surface sediments and comprise a large diversity of sulfate-reducing genera including *Desulfobacca, Desulfomonile, Desulfatiglans* and *Desulfatirhabdium*, as well as iron-reducing *Geobacter*. Interestingly, the sulfur-oxidizing epsilonproteobacterial genus *Sulfuricurvum* constitutes up to 5% of bacterial reads in the top 15 cm of sediment.

Besides *Proteobacteria*, dominant phyla in surface sediments include Parcubacteria (mainly unclassified and candidate groups Moranbacteria and Falkowbacteria), Chloroflexi (mainly classes Anaerolineae and Dehalococcoidia), Caldiserica (mainly genus *Caldiserica*), Aminicenantes, and Bacteroidetes (mainly vadinHA17, Sphingobacteriia, and unclassified). These members of Chloroflexi, Aminicenantes, and Bacteroidetes are widespread in freshwater sediments (Fiskal et al., 2019; Han et al., 2020) and groundwater (Farag et al., 2014). Caldiserica are often found in sulfidic environments, matching the thiosulfate-, sulfite-, and elemental sulfur-reducing metabolism of cultured isolates (Mori et al., 2009). Members of Parcubacteria, which account for 5-10% of reads in the top 35 cm and fall into the enigmatic Candidate Phyla Radiation (Brown et al., 2015), are widespread in diverse oxic and anoxic habitats, including lake water columns and terrestrial aquifers, and are potentially involved in carbohydrate metabolism (Wrighton et al., 2012; Nelson and Stegen, 2015; Vigneron et al., 2020). Besides Parcubacteria, significant percentages of Saccharibacteria, Zixibacteria (RBG-1), and Abscondibacteria (SR1; all up to ∼2%), and several other phyla affiliating with the Candidate Phyla Radiation (CPR), result in the CPR accounting for ∼10-18% of total bacterial sequence reads in the top 30 cm.

In the mid-column euxinic sediment, bacterial diversity is still high at the ZOTU level, ranging from 1800 ZOTUs in the upper 160 cm and decreasing to ∼800 below 700 cm. At the phylum level, Proteobacteria (of which almost all are Deltaproteobacteria), comprise between 8 and 15% of total bacterial sequences. Planctomycetes (mostly Phycisphaerae), Chloroflexi (mostly Dehalococcoidia groups GIF9, vadinBA26, MSBL5, and unclassified), Atribacterota, Aminicenantes, and Acidobacteria (mostly Subgroup 13) increase notably in relative abundances. None of these sequences have closely related cultured representatives. These phyla are typically detected in low-energy sedimentary environments in both the marine and freshwater systems (Hongxiang et al., 2008; Carr et al., 2015; Vuillemin et al., 2018; Han et al., 2020). Of note, peaks in Spirochaetaceae and Anaerolinaceae (class Anaerolineae within Chloroflexi) abundances coincide with the upper and lower sulfate depletion zone, as well as significant percentages of Parcubacteria (up to 4%; mostly Candidatus Magasanikbacteria) and Zixibacteria (up to 5%) throughout most of the mid-column interval.

The redox transition interval represents a transition not only in geochemical conditions but also in bacterial diversity. There is a small peak in diversity at the ZOTU level (957 OTUs) and a major shift in the dominant phyla, notably in the abundance of Aminicenantes sequences.

Compared to the overlying sediment, bacterial diversity in the deep glacial sediments is greatly reduced, with the total ZOTU numbers falling below 80 in parallel with the sharp decreasing cell numbers (Fig 3). Proteobacteria are again the most abundant phylum, with Alpha-, Beta- and Deltaproteobacteria together constituting 10-50% of total bacterial sequences there. Most of these are affiliated with the Rickettsiales (Alphaproteobacteria), Burkholderiales (Betaproteobacteria), Myxococcales, and Candidate clade Sva0485 (both Deltaproteobacteria), the latter of which has the genomic potential not only to reduce sulfate and oxidize sulfide, but also to cycle iron (Tan et al., 2019). Rhodobacterales (Alphaproteobacteria) exhibit a peak at 790 cm, within the deep sulfate depletion zone.

Actinobacteria, many of which are aerobic, are the second most abundant phylum in deep glacial sediments, constituting up to 78% of bacterial reads at some depths. Among the Acidobacteria, Subgroup 6 and Solibacteres are most abundant, with the notable presence of one ZOTU affiliated to the genus *Bryobacter*, an obligately aerobic chemo-organotrophic lineage of Solibacteres (Kulichevskaya et al., 2010), comprising ∼16% of bacterial reads at 890 cm depth. Firmicutes related to the classes Bacilli and Clostridia account for up to 8 and 15% of bacterial reads in glacial sediments, respectively, and have both been cited as keystone groups in the oligotrophic deep biosphere (Purkamo et al., 2016; Bose et al., 2020). While Parcubacteria make up 7% of bacterial reads in the glacial sediments, these ZOTUs belong to Candidatus Yanofskybacteria and Unclassified members, and are thus distinct from surface and mid-sediment Parcubacteria. Another noteworthy finding are the locally significant relative abundances of the CPR phyla Galena 15 (GAL15; 8.5%), Berkelbacteria (4%), Katanobacteria (WWE3; 9%) and Dependentiae (TM6; 5%). These phyla have characteristically small genomes that lack (known) core metabolic functions (Kantor et al., 2013; Hug et al., 2016). Their sub-micrometer cell size has earned them the name “nanobes” and allows them to be easily transported in groundwater (Luef et al., 2015). Altogether CPR phyla are 10-25% abundant in most glacial samples, which is in a similar range to sulfate-rich surface sediments, but higher than percentages in sulfate-depleted surface and mid-column sediments (1-7%)

### Phylum- and class-level variations in archaeal communities

A total of 380 archaeal ZOTUs (4,506,390 reads) were recovered, which could all be classified into a total of 13 different phyla (Fig. 6). Like Bacteria, Archaea are most diverse in surface sediment (Fig. 6), affiliating with seven different phyla of which Euryarchaeota of the candidate order Thermoprofundales (also known as Marine Benthic Group D; class Thermoplasmata; Supplementary Fig. 5) were most abundant (25-60% of archaeal reads). Genomic data on Thermoprofundales suggest protein fermentation as the dominant energy metabolism of this group (Lloyd et al., 2013; Zhou et al., 2019). Aenigmarchaeota, Bathyarchaeota, Diapherotrites, Pacearchaeota, and Woesearchaeota comprise most of the remainder of surface Archaea. Notably, Pacearchaeota have been reported to dominate surface waters of high-altitude oligotrophic lakes (Ortiz-Alvarez and Casamayor, 2016) and surface sediments of temperate lakes (Han et al., 2020). In mid-column sediments, the bathyarchaeotal Subgroup 5 (MCG-5) which genomic analyses have linked to energy metabolism via carbohydrate fermentation and acetogenesis (Zhou et al., 2018) are dominant. In the glacial sediments, archaeal diversity is reduced to 1-2 phyla per depth, though the sample from 790 cm depth should be interpreted with caution because only 7 reads were obtained as opposed to > 35000 reads from other glacial samples. In addition to Diapherotrites, aerobic Thaumarchaeaota affiliating with the ammonia-oxidizing *Nitrososphaera* were recovered from these deep sediments.

**Figure 6.**
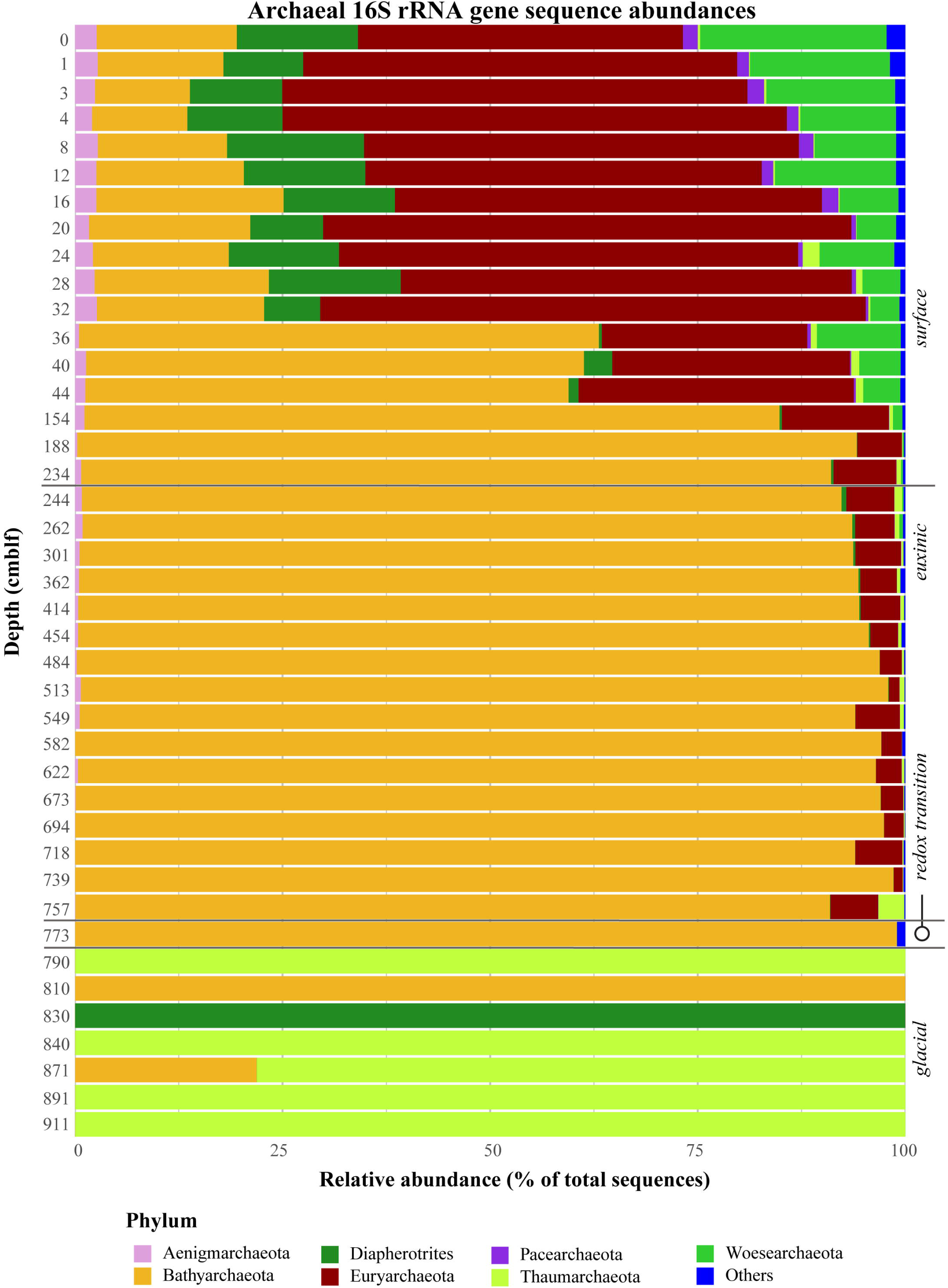
Relative abundances of Archaeal 16S rRNA gene sequences with depth. Sediment geological transitions are indicated on the right.

## DISCUSSION

Meromictic lakes are often cited as ancient ocean analogues and their sediments may retain geochemical signatures of past processes occurring in their anoxic waters. In this first complete biogeochemical characterization of the sediment record from a meromictic lake, we unveil both solid phase and porewater chemistry together with sedimentary microbial communities reflecting the first oxic, then suboxic, and finally anoxic ∼13.5 kyr history of the lake. Moreover, the well-preserved stratigraphy of our piston cores provides an environmental reconstruction of the history of Lake Cadagno.

Discrete geochemical periods resulted in the deposition of distinct sediment types, beginning with white powdery glacial till, rich in detrital clastic elements (Na, Al, K) and poor in iron sulfides, which was deposited ∼13.5-9.95 kyrs, prior to the establishment of meromixis and euxinia in Lake Cadagno. The late glacial sediments are low in carbonate content and TOC thus supporting sparse microbial communities. Interestingly, porewaters in these sediments contain relatively high concentrations of SO_4_^2-^, Mg^2+^, and Ca^2+^ (Fig. 2 & Supplementary Fig. 3B), which is a first indication that this deeply buried sediment might be influenced by subterranean aquifers. Sediment deposited during the anoxic-oxic transition period 9.95-9.01 kyrs (778-760 cm) is rich in Mn-, carbonate, and organic matter indicating a change in lake stratification conditions. The overlying 760 cm of euxinic sediment exhibits high accumulations of methane, DIC, and dissolved reduced metals resulting from the slow degradation of organic matter, except in very recent sediment dating to 220 years before present (40-0 cm) where steep chemical gradients indicate extremely rapid organic matter degradation and diagenesis.

In the following sections we present the relationships between dominant microbial communities and geochemistry in each of these sediment zones and individually discuss important microbial metabolisms involved in organic matter degradation including sulfur cycling, metal reduction, fermentation and methanogenesis. We also highlight the novel discovery of an oxidizing groundwater source generating a second redox transition zone in the deep subsurface with implications for deep sediment diagenesis.

### Extremely high microbial activity and rapid diagenesis in surface sediments

Most of the microbial activity in the Lake Cadagno sediments occurs within the top 20 cm where about 75% of the total organic carbon is remineralized (Fig. 2A) and the highest microbial cell numbers are observed (Fig. 3A). This very rapid degradation of organic matter suggests that TOC is highly reactive, which is not surprising given its mostly microbial and phytoplanktonic origin indicated by high N-content (Fig. 2A) and the short sinking depth of 20 m before deposition on the lake floor.

The presence of labile organic carbon in sulfate-rich surface sediments supports extremely diverse communities of microorganisms compared to euxinic sediments below 40 cm. These communities are remarkably similar to other lacustrine surface sediments despite the absence of an oxic-anoxic transition zone in Lake Cadagno sediments, with notably high abundances of Bacteroidetes, Verrucomicrobia, Alpha-, Beta-, and Gammaproteobacteria. In addition, the anoxic conditions in the water column and upper 20 cm favor Deltaproteobacteria and Chloroflexi, groups which are typically less common in surface sediments underlying oxic water. We also recovered significant percentages of sequences belonging to phyla that are ubiquitous, yet so far uncultivated. The CPR phyla constitute a large group of small-celled bacteria with unknown ecologies that have been postulated to be symbiotic based on their limited identifiable biosynthetic capabilities (Ferrari et al., 2014; Yeoh et al., 2016; Vigneron et al., 2020; Moreira et al., 2021). The abundance trends of CPR along with Chlamydiae, which are obligate symbionts of eukaryotic hosts (Dharamshi et al., 2020), suggests that they may simply be transported from the water column or surrounding soil and are selected against in the anoxic sediment, thus disappearing with depth. Eukaryotes such as protists may also survive in the deep oxidizing fluids which would thus explain the presence of highly abundant Rickettsiales which are also obligate intracellular bacteria (Perlman et al., 2006)

### Euxinic and redox transition sediment geochemistry and microbiology reflect depositional conditions

In the deeper euxinic sediment layers down to the glacial sediment, high concentrations of methane accumulate and organic matter becomes more recalcitrant. ^13^C-enriched TOC and higher C:N throughout most of the mid-column interval may reflect the selective removal of isotopically lighter, N-rich, and more labile organic matter of microbial and phytoplankton origin in shallow layers, leaving behind refractory terrestrial-dominated organic matter in deeper layers. Depth-profiles of Bacteria and Archaea reflect this trend as the mid-column sediments are dominated by clades specialized in the fermentative and acetogenic degradation of complex organic carbon substrates. These groups include Acidobacteria subgroup 13 which are possibly specialized in hemicellulose degradation (de Chaves et al., 2019), Anaerolineae members of which have been linked to protein and carbohydrate degradation and acetogenesis (Fullerton and Moyer, 2016; Xia et al., 2016; McIlroy et al., 2017), Dehalococcoidia which have been linked to reductive dehalogenation (Jugder et al., 2016), oxidation of various fatty acids, aromatic compounds, and even organo-sulphates (Wasmund et al., 2014), and carbohydrate degradation coupled to acetogenesis (Fincker et al., 2020), and Atribacterota and Aminicenantes which are abundant in methanogenic environments and have been linked to carbohydrate degradation (Farag et al., 2014; Kadnikov et al., 2019; Liu et al., 2019; Katayama et al., 2020). The Atribacterota in Lake Cadagno sediments exhibit very limited diversity with one dominant ZOTU comprising up to 28% of the total bacterial reads. Overall, carbon turnover at depth is likely extremely slow as reflected in the absence of a clear TOC depth gradient and the dominance of heterotrophic groups that are characteristic of energy-limited subsurface sediments such as Dehalococcoidia, Atribacterota, Aminicenantes, and Diapherotrites (Farag et al., 2014; Youssef et al., 2015; Nobu et al., 2016).

The Mn-rich layers just below (760-778 cm) correspond to the redox transition period from 9.95 to 9.01 kyrs. The sediment region corresponding to the onset of complete anoxia at 9.01 kyrs (760 cm) is characterized by a higher content of TOC (Fig. 2A), which was likely preserved due to decreased remineralization during anoxia, enhanced primary productivity due to warming and deglaciation. At that time, the activation of subaquatic springs combined with increased light availability (due to spring thawing) may have led to higher nutrient availability and primary productivity in the photic zone thus increasing the benthic oxygen demand leading to bottom water anoxia and enhancing organic carbon preservation (Wirth et al., 2013). The accompanying excursion to light δ^13^C values of -35 ‰, similar to the surface sediments, indicates material of pelagic origin (Bernasconi et al., 1997; Lehmann et al., 2002). The higher TOC content in the redox transition zone appears to support greater microbial cell numbers. Although only one sample was sequenced from within these transition sediments, the microbial diversity was strikingly different from the underlying late glacial sediments and the overlying mid-column sulfidic sediment (Fig. 4). This microbial community shift may be associated with differences in TOC quantity and quality, or the onset of oxidizing conditions not only at the time of deposition, but also in contemporary porewater gradients.

### Evidence for a deep, oxidizing source below the anoxic sediment column

Sediments situated below 778 cm depth are characterized by extremely low cell numbers (Fig. 3A), likely reflecting their low TOC content. Increasing ratios of Bacteria to Archaea in these sediments may reflect oxidizing conditions there (Chen et al., 2017; Deng et al., 2020). Similar sediment sequences of sulfidic sediment with underlying oxidized sediment can be found in euxinic basins such as the Black Sea recording a transition from oxic to anoxic conditions (Jørgensen et al., 2004). These ancient, deeply buried sediments can be subject to contemporary influences such as downwards-seeping sulfate in the Black Sea. In Lake Cadagno, sulfate is consumed in the upper 20 cm of sediment and the deep, ancient sediments are instead impacted by a deep source of sulfate. The upwards sulfate flux of about 5 µmol m^- 2^ d^-1^ overlaps with a downwards flux of methane of 87 µmol m^-2^ d^-1^ at around 770 cm depth. These fluxes are 1-2 orders of magnitude lower than at the surface depletion zone, partly due to the low porosity and permeability of clastic clays, and partly due to the decrease in reactivity of the organic matter available for microbial methanogenesis. The huge imbalance between electron donors and acceptors at the deep sulfate depletion zone indicates that other oxidants are available to consume methane, which does not penetrate into the glacial sediments. The presence of aerobic Thaumarchaeota for one suggest that O_2_ or nitrate are present, although the latter was measured and remained below detection limits.

Together, our biogeochemical data suggest the presence of an oxidizing groundwater source located at the base of the Lake Cadagno sediment column. The fact that free sulfide, Fe^2+^ and Mn^2+^ were not detectable below 820 cm suggests the presence of an oxidizing, sulfate-rich water source at the base of the sediment column, possibly containing molecular oxygen. This groundwater may either simply be ancient water trapped in pore spaces of the clay-rich glacial till similar to in Baltic Sea sediments (Jørgensen et al., 2020) or a subsurface aquifer which also feeds the underwater springs at 8-10 m water depth (Del Don et al., 2001). In the first scenario, a pool of Fe- and Mn-oxides of clastic origin may drive a cryptic sulfur cycle, chemically re-oxidizing sulfur compounds as previously suggested (Su et al., 2019). In the second scenario, groundwater flow may transport electron acceptors and even microbial cells into the deep sediments. The presence of Fe-oxides and oxidized sulfur intermediates like S^0^ (Fig. 2E), and the co-occurrence of microaerobic to facultatively aerobic taxa such as *Bryobacter* (Acidobacteria) and *Nitrosphaera* (Thaumarchaeota) (Fig. 3&4), suggests that this groundwater might even contain dissolved O_2_.

The presence of a deep, oxidizing groundwater source generates an inverse redox gradient and fresh source of electron acceptors deep within the sediment. Such an inverse gradient has also been reported in Pacific Ocean sediments (D’Hondt et al., 2004) and the Baltic Sea (Jørgensen et al., 2020) and may therefore be a widespread phenomenon in porous sedimentary systems. Although lower fluxes of methane and microbial cell numbers suggest that microbial activity is less than at the surface, deep sources of electron acceptors could contribute to the remineralization of organic matter sequestered deep below the surface.

### Surprising potential for sulfur cycling in the deep sediments of Lake Cadagno

Sulfur cycling appears to play an important role throughout the sediments of Lake Cadagno. Although the bottom waters of Lake Cadagno are permanently anoxic and no nitrate or nitrite was detected in porewaters, the presence of the sulfur-oxidizing *Sulfuricurvum* (5% of bacterial reads) in the upper 12 cm of sediments suggests either deposition from the water column or that intermittent oxygenation of bottom waters by density-driven currents drives oxidative sulfur cycling there. Sulfate reduction occurs prior to Mn- and Fe-reduction in the uppermost sediment layers and consumes sulfate completely within the top 20 cm. Abundances of known sulfate-reducing Deltaproteobacteria (*Desulfobacca, Desulfomonile, Desulfatiglans* and *Desulfatirhabdium*) coincide with the decreasing concentration of sulfate. Below the sulfate depletion zone, the abundance of potential sulfur-cycling taxa, in particular Caldiserica, in the mid-column sediment suggests that internal cycling of sulfur may be possible long after measurable sulfate and sulfur have disappeared (Wasmund et al., 2016; Anantharaman et al., 2018). This is consistent with previous findings of active microbial sulfur cycling in sulfate-depleted sediments (Pellerin et al., 2018) and would be worthy of future investigation.

The sulfate penetration depth is regulated by both sulfate concentration, organic carbon reactivity, and sediment deposition rates. In Lake Cadagno, the relatively shallow sulfate depletion zone is in part due to sulfate limitation, in contrast to euxinic basins such as the Black Sea and the Cariaco Basin, where the sulfate penetration depth is on the order of several meters (Werne et al., 2003; Jørgensen et al., 2004). Low organic matter deposition rates lead to a reduction in microbial activity and subsequent downward migration of the sulfate-methane transition zone (SMTZ). The relatively low sediment deposition rates (2-2.5 mm yr^-1^) in Lake Cadagno (Wirth et al., 2013; Bueche and Junier, 2016) compared to the Black Sea (50 mm yr^-1^; (Henkel et al., 2012)) and the Cariaco Basin (30 mm yr^-1^; (Werne et al., 2003)) do not fit this trend and may therefore depend on quality and not quantity of the organic matter supply.

In the Lake Cadagno sediment, methanogenesis is initiated regardless of the presence of sulfate and thus there is no true SMTZ. The strong overlaps in CH_4_ and SO_4_^2-^ concentration profiles, and the near-linear CH_4_ gradient, which results in methane depletion only at the sediment surface, suggest that most methane in the upper 25 cm is diffusing upward from deeper layers, and that CH_4_ is mainly consumed at the sediment surface and water column.

This linear methane gradient (Fig. 1E) along with the failure to detect key anaerobic methane-oxidizing archaea suggests that there is minimal anaerobic oxidation of methane (AOM) occurring in these sediments (Schubert et al., 2011). Instead, sulfate reducers appear to preferably degrade energy-rich fresh organic matter over energy-poor methane.

### Metal reduction, fermentation, and methanogenesis are involved in the slow remineralization of deeply buried carbon

It has previously been shown that Mn and Fe in the Lake Cadagno sediments do not exhibit any considerable correlation to other elements and are not attributable to specific lacustrine or terrestrial lithologies (Wirth et al., 2013). This is because these biogeochemically reactive metals are involved in redox and diagenetic recycling processes within sediments and pore waters, erasing any potential source-related connection to the other elements. Indeed, the accumulation of high concentrations of dissolved Mn^2+^ and Fe^2+^ in porewaters below 40 cm (Fig. 2D) suggest that microbial metal reduction is an important driver of organic carbon remineralization in the euxinic sediments of Lake Cadagno. Semi-quantitative XRF measurements of Fe and Mn indicate that total Fe is much more abundant than Mn yet the higher porewater concentrations of Mn^2+^ (Fig. 2D) suggest that it is preferentially reduced, possibly owing to its higher energetic yields.

While both Mn- and Fe-oxides may be abiotically reduced by sulfide in surface sediments (Fig. 2E), active microbial metal reduction is likely occurring in the sulfate-depleted deep sediments of Lake Cadagno. Despite being commonly isolated from diverse environments including the deep subsurface (Coates et al., 1996), we recovered only four ZOTUs (> 0.5% abundant) affiliating with the iron-reducing genus *Geobacter* (Desulfuromonadales). Most known sulfur/sulfate reducers are also facultative metal reducers and may thus indirectly or directly drive Fe- and Mn-oxide reduction. Nonetheless, sequences belonging to known facultative metal reducing families such as *Desulfobulbaceae, Desulfovibirionaceae*, and *Desulfobacteraceae* are mostly absent from the deep sediment. The phylogenetic range of prokaryotes known to reduce metals also includes fermentative microorganisms, such as *Clostridium* and *Bacillus*, capable of transferring electron equivalents to metal oxides and is constantly expanding, for example to the Bacteroidetes (Vandieken et al., 2018). The ubiquitous distribution of Firmicutes and Bacteroidetes sequences throughout the sediment column (2-3% of sequences) and peaks in abundances of 23% and 16%, respectively, at the redox interface around 800-840 cm suggest that they may be involved in metal redox cycling. Together, our data suggest that the intensely studied model metal-reducers such as *Geobacter* spp., are of relatively minor importance in these deep sediments compared to more versatile microorganisms, including fermenters and possibly unknown groups as has been shown for marine suboxic sediments (Reyes et al., 2016).

Fermentative metabolisms are not only tightly linked to metal cycling but also to methanogenesis. Bacterial groups possibly living in syntrophic association with methanogens such as Spirochaetes and Anaerolineae oxidizing acetate and degrading hydrocarbons, respectively, in concurrence with anaerobic methane production were detected at both the surface and deep sulfate depletion zones (Lee et al., 2015; Concheri et al., 2017). Atribacteria are abundant throughout the mid-column sediments (up to 28% of total bacterial reads) likely due to the high energy yields of sugar fermentation. They have also been suggested as H_2_-producing syntrophic partners for methanogens based on their ubiquitous presence in anoxic, methane-rich habitats (Carr et al., 2015; Lee et al., 2018). Bacterial phyla relying exclusively on fermentation for energy acquisition such as Parcubacteria, Bacteroidetes, Chloroflexi, and Aminicenantes (Hug et al., 2013; Farag et al., 2014; León Zayas et al., 2017) are also dominant throughout the sediment record.

Despite high methane concentrations in the Lake Cadagno sediments, methanogen populations detected by 16S rRNA gene sequencing represent a very minor fraction of the microbial community. This observation common to low-energy environments can be explained by small populations of methanogens subsisting at the biological energy quantum and producing small amounts of methane which accumulates to high concentrations over time (Hoehler et al., 2001; Orsi et al., 2020). We detected only one potential group of methanogens (Supplementary Fig. 5) and no methanotrophs of the ANME-2d class nor of AOM-associated archaea which have previously been detected in Lake Cadagno sediments (Schubert et al., 2011; Su et al., 2019), but other methane-cycling archaea may simply not have been covered by our 16S rRNA gene survey because they generally exhibit low abundances.

## ACKNOWLEDGMENTS

This study was supported by Swiss National Science Foundation (SNF) grant No. 182096 (M.A.L.). We thank the entire 2019 Cadagno sampling crew for assistance in the field, and especially the Alpine Biology Center Foundation (Switzerland) for use of its research facilities. We also acknowledge Iso Christl, Madalina Jaggi, Rachele Ossola, Annika Fiskal, and Clemens Glombitza for their support with chemical analyses along with Longhui Deng for support with bioinformatic analyses.

## REFERENCES

Anantharaman, K., Hausmann, B., Jungbluth, S. P., Kantor, R. S., Lavy, A., Warren, L. A., et al. (2018). Expanded diversity of microbial groups that shape the dissimilatory sulfur cycle. The ISME Journal 12, 1715–1728. doi:10.1038/s41396-018-0078-0.

Bak, F., and Pfennig, N. (1991). Microbial sulfate reduction in littoral sediment of Lake Constance. FEMS Microbiol Lett 85, 31–42. doi:10.1111/j.1574-6968.1991.tb04695.x.

Bernasconi, S. M., Barbieri, A., and Simona, M. (1997). Carbon and nitrogen isotope variations in sedimenting organic matter in Lake Lugano. Limnology and Oceanography 42, 1755–1765. doi:https://doi.org/10.4319/lo.1997.42.8.1755.

Blaauw, M., and Christen, J. A. (2011). Flexible paleoclimate age-depth models using an autoregressive gamma process. Bayesian Analysis 6, 457–474. doi:10.1214/11-BA618.

Blees, J., Niemann, H., Wenk, C. B., Zopfi, J., Schubert, C. J., Kirf, M. K., et al. (2014). Microaerobic bacterial methane oxidation in the chemocline and anoxic water column of deep south-Alpine Lake Lugano (Switzerland). Limnology and Oceanography 59, 311–324. doi:10.4319/lo.2014.59.2.0311.

Bose, H., Dutta, A., Roy, A., Gupta, A., Mukhopadhyay, S., Mohapatra, B., et al. (2020). Microbial diversity of drilling fluids from 3000&thinsp;m deep Koyna pilot borehole provides insights into the deep biosphere of continental earth crust. Scientific Drilling 27, 1–23. doi:https://doi.org/10.5194/sd-27-1-2020.

Boudreau, B. P. (1997). Diagenetic models and their implementation. Berlin: Springer.

Brown, C. T., Hug, L. A., Thomas, B. C., Sharon, I., Castelle, C. J., Singh, A., et al. (2015). Unusual biology across a group comprising more than 15% of domain Bacteria. Nature 523, 208–211. doi:10.1038/nature14486.

Bueche, M., and Junier, P. (2016). Effect of organic carbon and metal accumulation on the bacterial communities in sulphidogenic sediments. Environ Sci Pollut Res 23, 10443–10456. doi:10.1007/s11356-016-6056-z.

Burdige, D. J. (2007). Preservation of Organic Matter in Marine Sediments:L Controls, Mechanisms, and an Imbalance in Sediment Organic Carbon Budgets? Chem. Rev. 107, 467–485. doi:10.1021/cr050347q.

Cadillo□Quiroz, H., Bräuer, S., Yashiro, E., Sun, C., Yavitt, J., and Zinder, S. (2006). Vertical profiles of methanogenesis and methanogens in two contrasting acidic peatlands in central New York State, USA. Environmental Microbiology 8, 1428–1440. doi:10.1111/j.1462-2920.2006.01036.x.

Canfield, D. E. (1998). A new model for Proterozoic ocean chemistry. Nature 396, 450–453. doi:10.1038/24839.

Capone, D. G., and Kiene, R. P. (1988). Comparison of microbial dynamics in marine and freshwater sediments: Contrasts in anaerobic carbon catabolism1. Limnology and Oceanography 33, 725–749. doi:10.4319/lo.1988.33.4part2.0725.

Carlton, R. G., Walker, G. S., Klug, M. J., and Wetzel, R. G. (1989). Relative Values of Oxygen, Nitrate, and Sulfate to Terminal Microbial Processes in the Sediments of Lake Superior. Journal of Great Lakes Research 15, 133–140. doi:10.1016/S0380-1330(89)71467-2.

Carr, S. A., Orcutt, B. N., Mandernack, K. W., and Spear, J. R. (2015). Abundant Atribacteria in deep marine sediment from the Adélie Basin, Antarctica. Front. Microbiol. 6. doi:10.3389/fmicb.2015.00872.

Chen, J., Hanke, A., Tegetmeyer, H. E., Kattelmann, I., Sharma, R., Hamann, E., et al. (2017). Impacts of chemical gradients on microbial community structure. The ISME Journal 11, 920. doi:10.1038/ismej.2016.175.

Claesson, M. J., O’Sullivan, O., Wang, Q., Nikkilä, J., Marchesi, J. R., Smidt, H., et al. (2009). Comparative Analysis of Pyrosequencing and a Phylogenetic Microarray for Exploring Microbial Community Structures in the Human Distal Intestine. PLoS One 4. doi:10.1371/journal.pone.0006669.

Cline, J. D. (1969). Spectrophotometric determination of hydrogen sulfide in natural waters. Limnology and Oceanography 14, 454–458. doi:10.4319/lo.1969.14.3.0454.

Coates, J. D., Phillips, E. J., Lonergan, D. J., Jenter, H., and Lovley, D. R. (1996). Isolation of Geobacter species from diverse sedimentary environments. Appl. Environ. Microbiol. 62, 1531–1536.

Concheri, G., Stevanato, P., Zaccone, C., Shotyk, W., D’Orazio, V., Miano, T., et al. (2017). Rapid peat accumulation favours the occurrence of both fen and bog microbial communities within a Mediterranean, free-floating peat island. Sci Rep 7, 1–10. doi:10.1038/s41598-017-08662-y.

de Chaves, M. G., Silva, G. G. Z., Rossetto, R., Edwards, R. A., Tsai, S. M., and Navarrete, A. A. (2019). Acidobacteria Subgroups and Their Metabolic Potential for Carbon Degradation in Sugarcane Soil Amended With Vinasse and Nitrogen Fertilizers. Front. Microbiol. 10. doi:10.3389/fmicb.2019.01680.

Dean, W. E., and Gorham, E. (1998). Magnitude and significance of carbon burial in lakes, reservoirs, and peatlands. Geology 26, 535–538. doi:10.1130/0091-7613(1998)026<0535:MASOCB>2.3.CO;2.

Del Don, C., Hanselmann, K. W., Peduzzi, R., and Bachofen, R. (2001). The meromictic alpine Lake Cadagno: Orographical and biogeochemical description. Aquat. sci. 63, 70–90. doi:10.1007/PL00001345.

Deng, L., Bölsterli, D., Kristensen, E., Meile, C., Su, C.-C., Bernasconi, S. M., et al. (2020). Macrofaunal control of microbial community structure in continental margin sediments. PNAS 117, 15911–15922. doi:10.1073/pnas.1917494117.

Dharamshi, J. E., Tamarit, D., Eme, L., Stairs, C. W., Martijn, J., Homa, F., et al. (2020). Marine Sediments Illuminate Chlamydiae Diversity and Evolution. Current Biology 30, 1032–1048.e7. doi:10.1016/j.cub.2020.02.016.

D’Hondt, S., Jørgensen, B. B., Miller, D. J., Batzke, A., Blake, R., Cragg, B. A., et al. (2004). Distributions of Microbial Activities in Deep Subseafloor Sediments. Science 306, 2216–2221. doi:10.1126/science.1101155.

Farag, I. F., Davis, J. P., Youssef, N. H., and Elshahed, M. S. (2014). Global Patterns of Abundance, Diversity and Community Structure of the Aminicenantes (Candidate Phylum OP8). PLoS One 9. doi:10.1371/journal.pone.0092139.

Ferrari, B., Winsley, T., Ji, M., and Neilan, B. (2014). Insights into the distribution and abundance of the ubiquitous Candidatus Saccharibacteria phylum following tag pyrosequencing. Sci Rep 4, 3957. doi:10.1038/srep03957.

Fincker, M., Huber, J. A., Orphan, V. J., Rappé, M. S., Teske, A., and Spormann, A. M. (2020). Metabolic strategies of marine subseafloor Chloroflexi inferred from genome reconstructions. Environmental Microbiology 22, 3188–3204. doi:https://doi.org/10.1111/1462-2920.15061.

Fiskal, A., Deng, L., Michel, A., Eickenbusch, P., Han, X., Lagostina, L., et al. (2019). Effects of eutrophication on sedimentary organic carbon cycling in five temperate lakes. Biogeosciences 16, 3725–3746. doi:https://doi.org/10.5194/bg-16-3725-2019.

Fullerton, H., and Moyer, C. L. (2016). Comparative Single-Cell Genomics of Chloroflexi from the Okinawa Trough Deep-Subsurface Biosphere. Appl. Environ. Microbiol. 82, 3000–3008. doi:10.1128/AEM.00624-16.

Han, X., Schubert, C. J., Fiskal, A., Dubois, N., and Lever, M. A. (2020). Eutrophication as a driver of microbial community structure in lake sediments. Environmental Microbiology 22, 3446–3462. doi:https://doi.org/10.1111/1462-2920.15115.

Hansel, C. M., Lentini, C. J., Tang, Y., Johnston, D. T., Wankel, S. D., and Jardine, P. M. (2015). Dominance of sulfur-fueled iron oxide reduction in low-sulfate freshwater sediments. The ISME Journal 9, 2400–2412. doi:10.1038/ismej.2015.50.

Henkel, S., Mogollón, J. M., Nöthen, K., Franke, C., Bogus, K., Robin, E., et al. (2012). Diagenetic barium cycling in Black Sea sediments – A case study for anoxic marine environments. Geochimica et Cosmochimica Acta 88, 88–105. doi:10.1016/j.gca.2012.04.021.

Herlemann, D. P., Labrenz, M., Jürgens, K., Bertilsson, S., Waniek, J. J., and Andersson, A. F. (2011). Transitions in bacterial communities along the 2000 km salinity gradient of the Baltic Sea. The ISME Journal 5, 1571–1579. doi:10.1038/ismej.2011.41.

Hoehler, T. M., Alperin, M. J., Albert, D. B., and Martens, C. S. (2001). Apparent minimum free energy requirements for methanogenic Archaea and sulfate-reducing bacteria in an anoxic marine sediment. FEMS Microbiology Ecology 38, 33–41. doi:10.1111/j.1574-6941.2001.tb00879.x.

Hongxiang, X., Min, W., Xiaogu, W., Junyi, Y., and Chunsheng, W. (2008). Bacterial diversity in deep-sea sediment from northeastern Pacific Ocean. Acta Ecologica Sinica 28, 479–485. doi:10.1016/S1872-2032(08)60026-8.

Hug, L. A., Castelle, C. J., Wrighton, K. C., Thomas, B. C., Sharon, I., Frischkorn, K. R., et al. (2013). Community genomic analyses constrain the distribution of metabolic traits across the Chloroflexi phylum and indicate roles in sediment carbon cycling. Microbiome 1, 22. doi:10.1186/2049-2618-1-22.

Hug, L. A., Thomas, B. C., Sharon, I., Brown, C. T., Sharma, R., Hettich, R. L., et al. (2016). Critical biogeochemical functions in the subsurface are associated with bacteria from new phyla and little studied lineages. Environmental Microbiology 18, 159–173. doi:10.1111/1462-2920.12930.

Jørgensen, B. B., Andrén, T., and Marshall, I. P. G. (2020). Sub-seafloor biogeochemical processes and microbial life in the Baltic Sea. Environmental Microbiology 22, 1688–1706. doi:10.1111/1462-2920.14920.

Jørgensen, B. B., Böttcher, M. E., Lüschen, H., Neretin, L. N., and Volkov, I. I. (2004). Anaerobic methane oxidation and a deep H2S sink generate isotopically heavy sulfides in Black Sea sediments 1 1Associate editor: D. E. Canfield. Geochimica et Cosmochimica Acta 68, 2095–2118. doi:10.1016/j.gca.2003.07.017.

Jugder, B.-E., Ertan, H., Wong, Y. K., Braidy, N., Manefield, M., Marquis, C. P., et al. (2016). Genomic, transcriptomic and proteomic analyses of Dehalobacter UNSWDHB in response to chloroform. Environmental Microbiology Reports 8, 814–824. doi:https://doi.org/10.1111/1758-2229.12444.

Kadnikov, V., Mardanov, A., Beletsky, A., Karnachuk, O., and Ravin, N. (2019). Genome of the candidate phylum Aminicenantes bacterium from a deep subsurface thermal aquifer revealed its fermentative saccharolytic lifestyle. Extremophiles 23. doi:10.1007/s00792-018-01073-5.

Kallmeyer, J., Pockalny, R., Ram Adhikari, R., Smith, D. C., and D’Hondt, S. (2012). Global distribution of microbial abundance and biomass in subseafloor sediment. Proceedings of the National Academy of Science 109, 16213–16216. doi:10.1073/pnas.1203849109.

Kantor, R. S., Wrighton, K. C., Handley, K. M., Sharon, I., Hug, L. A., Castelle, C. J., et al. (2013). Small Genomes and Sparse Metabolisms of Sediment-Associated Bacteria from Four Candidate Phyla. mBio 4. doi:10.1128/mBio.00708-13.

Karstens, L., Asquith, M., Davin, S., Fair, D., Gregory, W. T., Wolfe, A. J., et al. (2019). Controlling for Contaminants in Low-Biomass 16S rRNA Gene Sequencing Experiments. mSystems 4. doi:10.1128/mSystems.00290-19.

Katayama, T., Nobu, M. K., Kusada, H., Meng, X.-Y., Hosogi, N., Uematsu, K., et al. (2020). Isolation of a member of the candidate phylum ‘Atribacteria’ reveals a unique cell membrane structure. Nature Communications 11, 6381. doi:10.1038/s41467-020-20149-5.

Kempers, A. J., and Kok, C. J. (1989). Re-examination of the determination of ammonium as the indophenol blue complex using salicylate. Analytica Chimica Acta 221, 147–155. doi:10.1016/S0003-2670(00)81948-0.

Kulichevskaya, I. S., Suzina, N. E., Liesack, W., and Dedysh, S. N. (2010). Bryobacter aggregatus gen. nov., sp. nov., a peat-inhabiting, aerobic chemo-organotroph from subdivision 3 of the Acidobacteria. International Journal of Systematic and Evolutionary Microbiology, 60, 301–306. doi:10.1099/ijs.0.013250-0.

LaRowe, D. E., Arndt, S., Bradley, J. A., Estes, E. R., Hoarfrost, A., Lang, S. Q., et al. (2020). The fate of organic carbon in marine sediments - New insights from recent data and analysis. Earth-Science Reviews 204, 103146. doi:10.1016/j.earscirev.2020.103146.

Lee, S.-H., Park, J.-H., Kim, S.-H., Yu, B. J., Yoon, J.-J., and Park, H.-D. (2015). Evidence of syntrophic acetate oxidation by Spirochaetes during anaerobic methane production. Bioresour Technol 190, 543–549. doi:10.1016/j.biortech.2015.02.066.

Lee, Y. M., Hwang, K., Lee, J. I., Kim, M., Hwang, C. Y., Noh, H.-J., et al. (2018). Genomic Insight Into the Predominance of Candidate Phylum Atribacteria JS1 Lineage in Marine Sediments. Front Microbiol 9. doi:10.3389/fmicb.2018.02909.

Lehmann, M. F., Bernasconi, S. M., Barbieri, A., and McKenzie, J. A. (2002). Preservation of organic matter and alteration of its carbon and nitrogen isotope composition during simulated and in situ early sedimentary diagenesis. Geochimica et Cosmochimica Acta 66, 3573–3584. doi:10.1016/S0016-7037(02)00968-7.

León□Zayas, R., Peoples, L., Biddle, J. F., Podell, S., Novotny, M., Cameron, J., et al. (2017). The metabolic potential of the single cell genomes obtained from the Challenger Deep, Mariana Trench within the candidate superphylum Parcubacteria (OD1). Environmental Microbiology 19, 2769–2784. doi:10.1111/1462-2920.13789.

Lever, M. A., Torti, A., Eickenbusch, P., Michaud, A. B., Šantl-Temkiv, T., and Jørgensen, B. B. (2015). A modular method for the extraction of DNA and RNA, and the separation of DNA pools from diverse environmental sample types. Front. Microbiol. 6. doi:10.3389/fmicb.2015.00476.

Liu, Y.-F., Qi, Z.-Z., Shou, L.-B., Liu, J.-F., Yang, S.-Z., Gu, J.-D., et al. (2019). Anaerobic hydrocarbon degradation in candidate phylum ‘Atribacteria’ (JS1) inferred from genomics. The ISME Journal 13, 2377–2390. doi:10.1038/s41396-019-0448-2.

Lloyd, K. G., Schreiber, L., Petersen, D. G., Kjeldsen, K. U., Lever, M. A., Steen, A. D., et al. (2013). Predominant archaea in marine sediments degrade detrital proteins. Nature 496, 215–218. doi:10.1038/nature12033.

Lovley, D. R., and Phillips, E. J. P. (1986). Organic Matter Mineralization with Reduction of Ferric Iron in Anaerobic Sediments. Appl. Environ. Microbiol. 51, 683–689.

Luef, B., Frischkorn, K. R., Wrighton, K. C., Holman, H.-Y. N., Birarda, G., Thomas, B. C., et al. (2015). Diverse uncultivated ultra-small bacterial cells in groundwater. Nature Communications 6, 6372. doi:10.1038/ncomms7372.

McIlroy, S. J., Kirkegaard, R. H., Dueholm, M. S., Fernando, E., Karst, S. M., Albertsen, M., et al. (2017). Culture-Independent Analyses Reveal Novel Anaerolineaceae as Abundant Primary Fermenters in Anaerobic Digesters Treating Waste Activated Sludge. Front. Microbiol. 8. doi:10.3389/fmicb.2017.01134.

Moreira, D., Zivanovic, Y., López-Archilla, A. I., Iniesto, M., and López-García, P. (2021). Reductive evolution and unique predatory mode in the CPR bacterium Vampirococcus lugosii. Nat Commun 12, 2454. doi:10.1038/s41467-021-22762-4.

Mori, K., Yamaguchi, K., Sakiyama, Y., Urabe, T., and Suzuki, K. (2009). Caldisericum exile gen. nov., sp. nov., an anaerobic, thermophilic, filamentous bacterium of a novel bacterial phylum, Caldiserica phyl. nov., originally called the candidate phylum OP5, and description of Caldisericaceae fam. nov., Caldisericales ord. nov. and Caldisericia classis nov. International Journal of Systematic and Evolutionary Microbiology, 59, 2894–2898. doi:10.1099/ijs.0.010033-0.

Nelson, W. C., and Stegen, J. C. (2015). The reduced genomes of Parcubacteria (OD1) contain signatures of a symbiotic lifestyle. Front. Microbiol. 6. doi:10.3389/fmicb.2015.00713.

Nobu, M. K., Dodsworth, J. A., Murugapiran, S. K., Rinke, C., Gies, E. A., Webster, G., et al. (2016). Phylogeny and physiology of candidate phylum ‘Atribacteria’ (OP9/JS1) inferred from cultivation-independent genomics. The ISME Journal 10, 273–286. doi:10.1038/ismej.2015.97.

Ohkuma, M., and Kudo, T. (1998). Phylogenetic analysis of the symbiotic intestinal microflora of the termite Cryptotermes domesticus. FEMS Microbiol Lett 164, 389–395. doi:10.1111/j.1574-6968.1998.tb13114.x.

Onstott, T. C., Phelps, T. J., Kieft, T., Colwell, F. S., Balkwill, D. L., Fredrickson, J. K., et al. (1999). “A Global Perspective on the Microbial Abundance and Activity in the Deep Subsurface,” in Enigmatic Microorganisms and Life in Extreme Environments Cellular Origin and Life in Extreme Habitats., ed. J. Seckbach (Dordrecht: Springer Netherlands), 487–500. doi:10.1007/978-94-011-4838-2_38.

Orsi, W. D., Schink, B., Buckel, W., and Martin, W. F. (2020). Physiological limits to life in anoxic subseafloor sediment. FEMS Microbiology Reviews 44, 219–231. doi:10.1093/femsre/fuaa004.

Ortiz-Alvarez, R., and Casamayor, E. O. (2016). High occurrence of Pacearchaeota and Woesearchaeota (Archaea superphylum DPANN) in the surface waters of oligotrophic high-altitude lakes. Environ Microbiol Rep 8, 210–217. doi:10.1111/1758-2229.12370.

Parkes, R. J., Webster, G., Cragg, B. A., Weightman, A. J., Newberry, C. J., Ferdelman, T. G., et al. (2005). Deep sub-seafloor prokaryotes stimulated at interfaces over geological time. Nature 436, 390–394. doi:10.1038/nature03796.

Pellerin, A., Antler, G., Røy, H., Findlay, A., Beulig, F., Scholze, C., et al. (2018). The sulfur cycle below the sulfate-methane transition of marine sediments. Geochimica et Cosmochimica Acta 239, 74–89. doi:10.1016/j.gca.2018.07.027.

Perlman, S. J., Hunter, M. S., and Zchori-Fein, E. (2006). The emerging diversity of Rickettsia. Proceedings of the Royal Society B: Biological Sciences 273, 2097–2106. doi:10.1098/rspb.2006.3541.

Poulton, S. W., Fralick, P. W., and Canfield, D. E. (2004). The transition to a sulphidic ocean ∼ 1.84 billion years ago. Nature 431, 173–177. doi:10.1038/nature02912.

Purkamo, L., Bomberg, M., Kietäväinen, R., Salavirta, H., Nyyssönen, M., Nuppunen-Puputti, M., et al. (2016). Microbial co-occurrence patterns in deep Precambrian bedrock fracture fluids. Biogeosciences 13, 3091–3108. doi:https://doi.org/10.5194/bg-13-3091-2016.

Ravasi, D. F., Peduzzi, S., Guidi, V., Peduzzi, R., Wirth, S. B., Gilli, A., et al. (2012). Development of a real-time PCR method for the detection of fossil 16S rDNA fragments of phototrophic sulfur bacteria in the sediments of Lake Cadagno. Geobiology 10, 196–204. doi:10.1111/j.1472-4669.2012.00326.x.

Reimer, P. J., Bard, E., Bayliss, A., Beck, J. W., Blackwell, P. G., Ramsey, C. B., et al. (2013). IntCal13 and Marine13 Radiocarbon Age Calibration Curves 0–50,000 Years cal BP. Radiocarbon 55, 1869–1887. doi:10.2458/azu_js_rc.55.16947.

Reyes, C., Dellwig, O., Dähnke, K., Gehre, M., Noriega-Ortega, B. E., Böttcher, M. E., et al. (2016). Bacterial communities potentially involved in iron-cycling in Baltic Sea and North Sea sediments revealed by pyrosequencing. FEMS Microbiology Ecology 92. doi:10.1093/femsec/fiw054.

Roden, E. E., and Wetzel, R. G. (1996). Organic carbon oxidation and suppression of methane production by microbial Fe(III) oxide reduction in vegetated and unvegetated freshwater wetland sediments. Limnology and Oceanography 41, 1733–1748. doi:10.4319/lo.1996.41.8.1733.

Schink, B. (1997). Energetics of syntrophic cooperation in methanogenic degradation. Microbiol. Mol. Biol. Rev. 61, 262–280.

Schubert, C. J., Vazquez, F., Lösekann-Behrens, T., Knittel, K., Tonolla, M., and Boetius, A. (2011). Evidence for anaerobic oxidation of methane in sediments of a freshwater system (Lago di Cadagno). FEMS Microbiol Ecol 76, 26–38. doi:10.1111/j.1574-6941.2010.01036.x.

Sheik, C. S., Reese, B. K., Twing, K. I., Sylvan, J. B., Grim, S. L., Schrenk, M. O., et al. (2018). Identification and Removal of Contaminant Sequences From Ribosomal Gene Databases: Lessons From the Census of Deep Life. Front. Microbiol. 9. doi:10.3389/fmicb.2018.00840.

Sørensen, K. B., and Teske, A. (2006). Stratified Communities of Active Archaea in Deep Marine Subsurface Sediments. Appl. Environ. Microbiol. 72, 4596–4603. doi:10.1128/AEM.00562-06.

Stookey, L. (1970). Ferrozine---a new spectrophotometric reagent for iron. Analytical Chemistry 42, 779–781.

Su, G., Zopfi, J., Yao, H., Steinle, L., Niemann, H., and Lehmann, M. F. (2019). Manganese/iron-supported sulfate-dependent anaerobic oxidation of methane by archaea in lake sediments. Limnology and Oceanography n/a. doi:10.1002/lno.11354.

Tan, S., Liu, J., Fang, Y., Hedlund, B. P., Lian, Z.-H., Huang, L.-Y., et al. (2019). Insights into ecological role of a new deltaproteobacterial order Candidatus Acidulodesulfobacterales by metagenomics and metatranscriptomics. The ISME Journal 13, 2044–2057. doi:10.1038/s41396-019-0415-y.

Thomas, C., Francke, A., Vogel, H., Wagner, B., and Ariztegui, D. (2020). Weak influence of paleoenvironmental conditions on the subsurface biosphere of lake Ohrid in the last 515 ka. Earth ArXiv doi:10.31223/osf.io/jmye6.

Urban, N. R., Brezonik, P. L., Baker, L. A., and Sherman, L. A. (1994). Sulfate reduction and diffusion in sediments of Little Rock Lake, Wisconsin. Limnology and Oceanography 39, 797–815. doi:10.4319/lo.1994.39.4.0797.

Vandieken, V., Marshall, I. P. G., Niemann, H., Engelen, B., and Cypionka, H. (2018). Labilibaculum manganireducens gen. nov., sp. nov. and Labilibaculum filiforme sp. nov., Novel Bacteroidetes Isolated from Subsurface Sediments of the Baltic Sea. Front. Microbiol. 8. doi:10.3389/fmicb.2017.02614.

Vigneron, A., Cruaud, P., Langlois, V., Lovejoy, C., Culley, A. I., and Vincent, W. F. (2020). Ultrasmall and abundant: Candidate phyla radiation bacteria are potential catalysts of carbon transformation in a thermokarst lake ecosystem. Limnology and Oceanography Letters 5, 212–220. doi:https://doi.org/10.1002/lol2.10132.

Vuillemin, A., and Ariztegui, D. (2013). Geomicrobiological investigations in subsaline maar lake sediments over the last 1500 years. Quaternary Science Reviews 71, 119–130. doi:10.1016/j.quascirev.2012.04.011.

Vuillemin, A., Ariztegui, D., Horn, F., Kallmeyer, J., and Orsi, W. D. (2018). Microbial community composition along a 50 000-year lacustrine sediment sequence. FEMS Microbiol Ecol 94. doi:10.1093/femsec/fiy029.

Wang, Y., and Qian, P.-Y. (2009). Conservative Fragments in Bacterial 16S rRNA Genes and Primer Design for 16S Ribosomal DNA Amplicons in Metagenomic Studies. PLoS One 4. doi:10.1371/journal.pone.0007401.

Wasmund, K., Cooper, M., Schreiber, L., Lloyd, K. G., Baker, B. J., Petersen, D. G., et al. (2016). Single-Cell Genome and Group-Specific dsrAB Sequencing Implicate Marine Members of the Class Dehalococcoidia (Phylum Chloroflexi) in Sulfur Cycling. mBio 7. doi:10.1128/mBio.00266-16.

Wasmund, K., Schreiber, L., Lloyd, K. G., Petersen, D. G., Schramm, A., Stepanauskas, R., et al. (2014). Genome sequencing of a single cell of the widely distributed marine subsurface Dehalococcoidia, phylum Chloroflexi. The ISME Journal 8, 383–397. doi:10.1038/ismej.2013.143.

Werne, J. P., Lyons, T. W., Hollander, D. J., Formolo, M. J., and Sinninghe Damsté, J. S. (2003). Reduced sulfur in euxinic sediments of the Cariaco Basin: sulfur isotope constraints on organic sulfur formation. Chemical Geology 195, 159–179. doi:10.1016/S0009-2541(02)00393-5.

Wirth, S. B., Gilli, A., Niemann, H., Dahl, T. W., Ravasi, D., Sax, N., et al. (2013). Combining sedimentological, trace metal (Mn, Mo) and molecular evidence for reconstructing past water-column redox conditions: The example of meromictic Lake Cadagno (Swiss Alps). Geochimica et Cosmochimica Acta 120, 220–238. doi:10.1016/j.gca.2013.06.017.

Wrighton, K. C., Thomas, B. C., Sharon, I., Miller, C. S., Castelle, C. J., VerBerkmoes, N. C., et al. (2012). Fermentation, Hydrogen, and Sulfur Metabolism in Multiple Uncultivated Bacterial Phyla. Science 337, 1661–1665. doi:10.1126/science.1224041.

Xia, Y., Wang, Y., Wang, Y., Chin, F. Y. L., and Zhang, T. (2016). Cellular adhesiveness and cellulolytic capacity in Anaerolineae revealed by omics-based genome interpretation. Biotechnology for Biofuels 9, 111. doi:10.1186/s13068-016-0524-z.

Xinguo, H., Schubert, C. J., Fiskal, A., Dubois, N., and Lever, M. A. Eutrophication as a driver of microbial community structure in lake sediments. Available at: https://europepmc.org/article/med/32510812 [Accessed June 14, 2020].

Yeoh, Y. K., Sekiguchi, Y., Parks, D. H., and Hugenholtz, P. (2016). Comparative Genomics of Candidate Phylum TM6 Suggests That Parasitism Is Widespread and Ancestral in This Lineage. Mol Biol Evol 33, 915–927. doi:10.1093/molbev/msv281.

Youssef, N. H., Rinke, C., Stepanauskas, R., Farag, I., Woyke, T., and Elshahed, M. S. (2015). Insights into the metabolism, lifestyle and putative evolutionary history of the novel archaeal phylum “Diapherotrites.” ISME J 9, 447–460. doi:10.1038/ismej.2014.141.

Yu, Y., Lee, C., Kim, J., and Hwang, S. (2005). Group-specific primer and probe sets to detect methanogenic communities using quantitative real-time polymerase chain reaction. Biotechnology and Bioengineering 89, 670–679. doi:10.1002/bit.20347.

Zhou, Z., Liu, Y., Lloyd, K. G., Pan, J., Yang, Y., Gu, J.-D., et al. (2019). Genomic and transcriptomic insights into the ecology and metabolism of benthic archaeal cosmopolitan, Thermoprofundales (MBG-D archaea). The ISME Journal 13, 885–901. doi:10.1038/s41396-018-0321-8.

Zhou, Z., Pan, J., Wang, F., Gu, J.-D., and Li, M. (2018). Bathyarchaeota: globally distributed metabolic generalists in anoxic environments. FEMS Microbiol Rev 42, 639–655. doi:10.1093/femsre/fuy023.

